# A promoter-proximal silencer modifies the activity of a shared enhancer to mediate divergent expression of *nub* and *pdm2* paralogs in wing development

**DOI:** 10.1101/2022.03.01.482523

**Authors:** Ryan Loker, Richard S. Mann

## Abstract

Duplication of genes and their associated *cis*-regulatory elements, or enhancers, is a key contributor to genome evolution and biological complexity. Moreover, many paralogs, particularly tandem duplicates, are fixed for long periods of time under the control of shared enhancers. However, in most cases the mechanism by which gene expression and function diverge following duplication is not known. Here we dissect the regulation and function of the paralogous *nubbin*/*pdm2* genes during wing development in *Drosophila melanogaster*. We show that these paralogs play a redundant role in the wing and that their expression relies on a single shared wing enhancer. However, the two genes differ in their ability to respond to this enhancer, with *nub* responding in all wing progenitor cells and *pdm2* only in a small subset. This divergence is a result of a *pdm2*-specific silencer element at the *pdm2* promoter that receives repressive input from the transcription factor Rotund. Repression through this silencer also depends on *nub*, allowing *pdm2* to fully respond to the wing enhancer when *nub* expression is perturbed and functional compensation to occur. Thus, expression divergence downstream of a shared enhancer arises as a consequence of silencing the promoter of one paralog.

## Introduction

A major driver of genome evolution and biological diversity is the acquisition of new genes via duplication (Ohno, 1970). As such, the mechanisms by which duplicated genes are retained in the genome and their potential evolutionary trajectories are of great interest. While the majority of genes resulting from duplication events are quickly lost through a process known as nonfunctionalization (Force et al., 1999), preservation of paralogs in the genome is thought to occur via a number of mechanisms (Lynch et al., 2001; Lynch & Force, 2000). Duplicated genes can diverge either by acquiring new functions (neofunctionalization (Ohno, 1970)) or by the partitioning of ancestral functions, as originally described by the Duplication-Degeneration-Complementation (DDC) or subfunctionalization models (Force et al., 1999)). However, preserved duplicates can also persist without an observable divergence in expression or function, where they may maintain functional redundancy over long periods of time (Kuzmin et al., 2020; Lan & Pritchard, 2016). Redundancy due to gene duplication has also been proposed to aid in robustness of biological processes in response to environmental or genetic perturbations (Gu et al., 2003; Osterwalder et al., 2018). In all eukaryotes that have been studied, a particularly strong propensity for co-expression is found for paralogs that exist close to one another in the genome, called tandem duplicates (Lan & Pritchard, 2016; Quintero-Cadena & Sternberg, 2016; Williams & Bowles, 2004). One explanation that has been proposed for the high prevalence of co-expression for tandem duplicates is that they may be under control of a shared set of *cis*-regulatory elements or enhancers. Co-dependence on a single regulatory element would also explain why some tandem duplicates have conserved synteny over long evolutionary time scales. However, the functional validation of shared enhancers has only been demonstrated in a small number of instances (Baudouin-Gonzalez et al., 2017; Lan & Pritchard, 2016). Moreover, because of their dependency on the same enhancers, it is less clear how tandem duplicates can evolve complementary or novel functions.

Here we present an analysis of the expression and function of the paralogous *nubbin* (*nub*, also known as *pdm1*) and *pdm2* genes during development of the wing appendage in *Drosophila melanogaster. nub* and *pdm2* arose in an ancient duplication near the base of the divergence of the Brachycera suborder around 200 million years ago (Figure 1A). Both genes encode similar POU-type homeodomain transcription factors (TFs) (Figure S1) and play a role in development of a wide variety of cell types (Cifuentes & García-Bellido, 1997; Corty et al., 2016; Ng et al., 1995; Yeo et al., 1995). In winged insects (Pterygota) *nub* is a conserved marker for wings and has been shown to be essential for wing development from beetles to flies (Averof & Cohen, 1997; Cifuentes & García-Bellido, 1997; Ng et al., 1995; Tomoyasu et al., 2009). *nub* is expressed throughout the wing primordia in the *Drosophila* wing imaginal disc, including the progenitors of both the distal hinge and wing blade. Strong wing phenotypes are observed in multiple alleles attributed to loss of *nub* expression (Cifuentes & García-Bellido, 1997; Ng et al., 1995). We show, however, that *nub* is dispensable for wing development in *Drosophila* as a result of compensation by *pdm2*, revealing a previously unknown redundancy of these paralogs during wing development. Further, the wing phenotype of a classical *nub* allele is a result of a deletion of a shared wing enhancer that is essential for both *nub* and *pdm2* expression in wing progenitor cells. Surprisingly, although the expression of both genes in the wing depend on the same enhancer, the expression patterns of *nub* and *pdm2* differ under normal developmental conditions. The cause of this divergent expression is a *pdm2*-specific silencer element localized to its promoter (∼20 kb from the enhancer) that blocks the response to the shared enhancer in most wing cells. Repression through this element requires direct input from the TF Rotund (Rn). *nub* is also required for *pdm2* repression and, in the absence of *nub, pdm2* responds to the shared enhancer in all wing cells, revealing the mechanism for its compensatory function. Altogether this study highlights a case in which redundant paralogs that depend on the same enhancer have divergent expression patterns because of repressive inputs into one of their promoters, thus revealing a novel mechanism for how tandem duplicates can evolve distinct functions. Finally, by characterizing the chromatin landscape of appendage cells in multiple conditions, we also reveal the widespread downstream function of *nub* and *pdm2* as transcriptional repressors.

**Figure 1.**
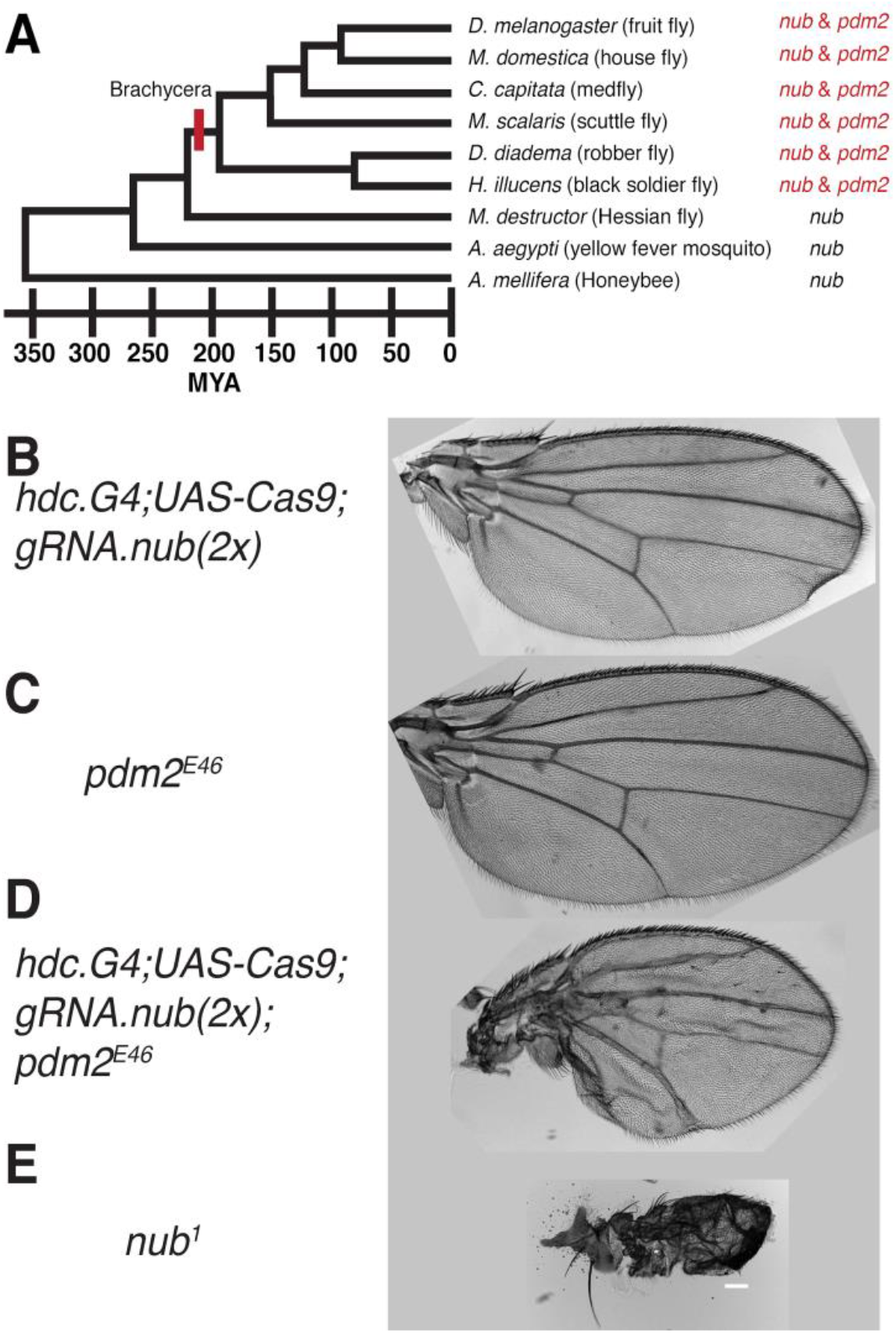
Functional redundancy of *nubbin* and *pdm2* during wing development. (**A**) Duplication of an ancestral proto *nub*/*pdm2* gene occurred in the Dipteran lineage near the base of divergence within the Brachycera suborder. Time span shown below is approximate millions of years ago (MYA). (**B-E**) Adult wing morphology with the following conditions: (**B**) Somatic CRISPR targeting *nubbin* coding exons with two gRNAs (**C**) *pdm2* truncated allele: *pdm2*^*E46*^ (**D**) Somatic CRISPR targeting *nubbin* coding exons with two gRNAs in a *pdm2*^*46*^ background. The resulting phenotype is a partial disruption of wing morphology due to the mosaic nature of somatic CRISPR (see Figure S2B). (**E**) *nub*^*1*^ allele. Scale bar is 100um.

## Results

### *Functional redundancy of* nub *and* pdm2 *in wing development*

Classical regulatory alleles of *nub* cause severe growth and tissue organization defects of the adult wing (Cifuentes & García-Bellido, 1997; Ng et al., 1995). Surprisingly, however, when Nubbin protein is specifically depleted during wing development, either through RNAi knockdown (López-Varea et al., 2021) or somatic CRISPR targeting of the gene locus (Figure 1C), the wings appear normal. To examine if the lack of wing phenotype could be due to a redundant function of the paralogous *pdm2* gene, we repeated the somatic CRISPR loss-of-function experiments in the background of the *pdm2*^*E46*^ allele, which produces a truncated protein as a result of a premature termination codon prior to the homeodomain (Yeo et al., 1995). Animals lacking *pdm2* are viable and have morphologically normal wings (Figure 1D). Furthermore, wing imaginal discs from *pdm2*^*E46*^ larvae lack staining by Pdm2-specific antibody (Figure S2A). When *nub* is mosaically removed in wings that also lack *pdm2* the adult appendage shows defects similar to wings containing clones of cells with classical *nub* regulatory alleles, characterized by reduction of hinge organization, overall wing size, and ectopic margin bristles in the interior blade region (Figure 1E, (Cifuentes & García-Bellido, 1997; Ng et al., 1995). The mosaic phenotype in this background is a result of incomplete knockout from the CRISPR targeting, which can be visualized from clonal loss of Nub protein in the wing disc (Figure S2B). These observations imply that 1) *pdm2* and *nub* can function redundantly during wing development and 2) that classic loss of function *nub* alleles disrupt the expression of both *nub* and *pdm2*. Below, we test and confirm both of these predictions.

### *Compensation of* nub *loss-of-function by upregulation of* pdm2

We next examined the expression of Nub and Pdm2 within wing progenitor tissue during development. Although Nub expression levels appear uniform throughout the appendage-generating domain of the wing imaginal disc, Pdm2 expression is limited to a subset of progenitor cells at the periphery of the appendage domain that express high levels, while more distal cells have weak or no detectable protein (Figure 2A-B). Notably, the anti-Nubbin antibody used here and previously (Averof & Cohen, 1997) recognizes both Nub and Pdm2 proteins while the anti-Pdm2 antibody specifically labels Pdm2 (data not shown). Given that Pdm2 is only appreciably expressed in the outer edge of the distal hinge domain, we conclude that anti-Nub staining in the central appendage domain is largely the result of Nub protein only, whereas in the edge of the distal hinge the signal results from the co-expression of Nub and Pdm2 (Figure 2C).

**Figure 2.**
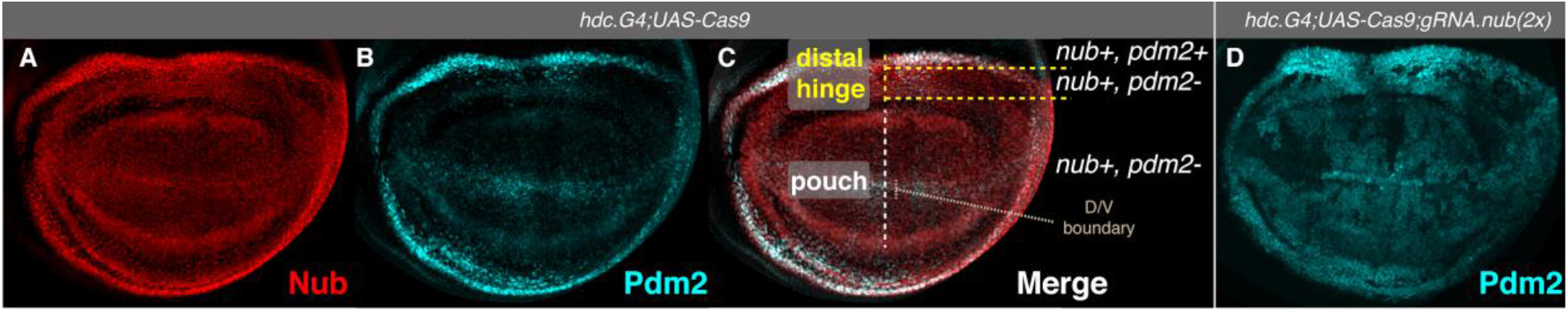
Expression of Nub and Pdm2 in the developing wing in normal and *nub* loss-of-function conditions. (**A**) Expression of Nub protein in the wing imaginal disc. Note that the anti-Nub antibody recognizes both Nub and Pdm2 (**B**) Expression of Pdm2 in the wing imaginal disc. High levels of Pdm2 protein are limited to the peripheral cells of the appendage domain.(**C**) Merged Nub/Pdm2 expression with relevant domains indicated. (**D**) Expression of Pdm2 in the wing imaginal disc upon targeting of *nub* with somatic CRISPR. Pdm2 levels are elevated in the majority of cells within the central domain of the appendage domain. Patches of cells with lower Pdm2 levels can be observed that are likely a result of incomplete Nub knockout, as the somatic CRISPR produces a mosaic loss-of-function (see Figure S2B).

The differences in expression between Nub and Pdm2 raise the question of how *pdm2* can compensate for loss of *nub* during wing development. We tested the idea that *pdm2* is upregulated in the absence of *nub*. Consistent with this idea, we find that Pdm2 levels increase throughout the wing pouch domain following loss of *nub* induced by somatic CRISPR (Figure 2D) or upon RNAi induced knockdown specifically in the Posterior compartment (Figure S3).

### *The* nub^1^ *allele is a regulatory mutant that removes both* nub *and* pdm2 *expression*

The results explained thus far demonstrate that both *nub* and *pdm2* are each sufficient for wing development, and therefore raise the question of how previously described *nub* alleles display strong wing phenotypes. Multiple viable alleles of *nub* have been described that develop wing malformations and each is associated with insertions or re-arrangements within non-coding regions near the *nub* locus (Ng et al., 1995). We focused on the *nub*^*1*^ allele, which displays a very strong wing phenotype (Figure 1E) (Cifuentes & García-Bellido, 1997; Ng et al., 1995). We first performed RNA-seq using wing discs from *nub*^*1*^ animals and compared the *nub* and *pdm2* transcript levels with wild type tissue. This analysis showed that *nub* levels were drastically reduced and *pdm2* transcripts were undetectable in the *nub*^*1*^ background (Figure 3A). Antibody staining of *nub*^*1*^ wing discs revealed that most of the tissue lacks Nubbin protein, with only two small clusters of cells retaining some expression, as previously reported (Cifuentes & García-Bellido, 1997). Antibody staining for Pdm2 revealed that it was nearly absent in *nub*^*1*^ wing discs, consistent with the RNA-seq analysis (Figure 3B). Analysis of mitotic clones with the *nub*^*1*^ allele reveals that cells in the domain that retain some Nub expression display lower levels of protein compared to wild type cells (Figure S4A). These results suggest that the phenotype associated with the *nub*^*1*^ allele is due to loss of expression of both *nub* and *pdm2* during wing development. To further establish that *nub*-associated wing phenotypes are associated with loss of both paralogs, we examined protein levels in wing discs of a different allele that causes a strong wing phenotype, *nub*^*2*^ (Ng et al., 1995). Consistent with the *nub*^*1*^ result, *nub*^*2*^ wing discs show a similar loss of Nub and Pdm2 (Figure S4B). In the next sections we investigate the mechanism by which expression of both genes is impaired in the *nub*^*1*^ genetic background.

**Figure 3.**
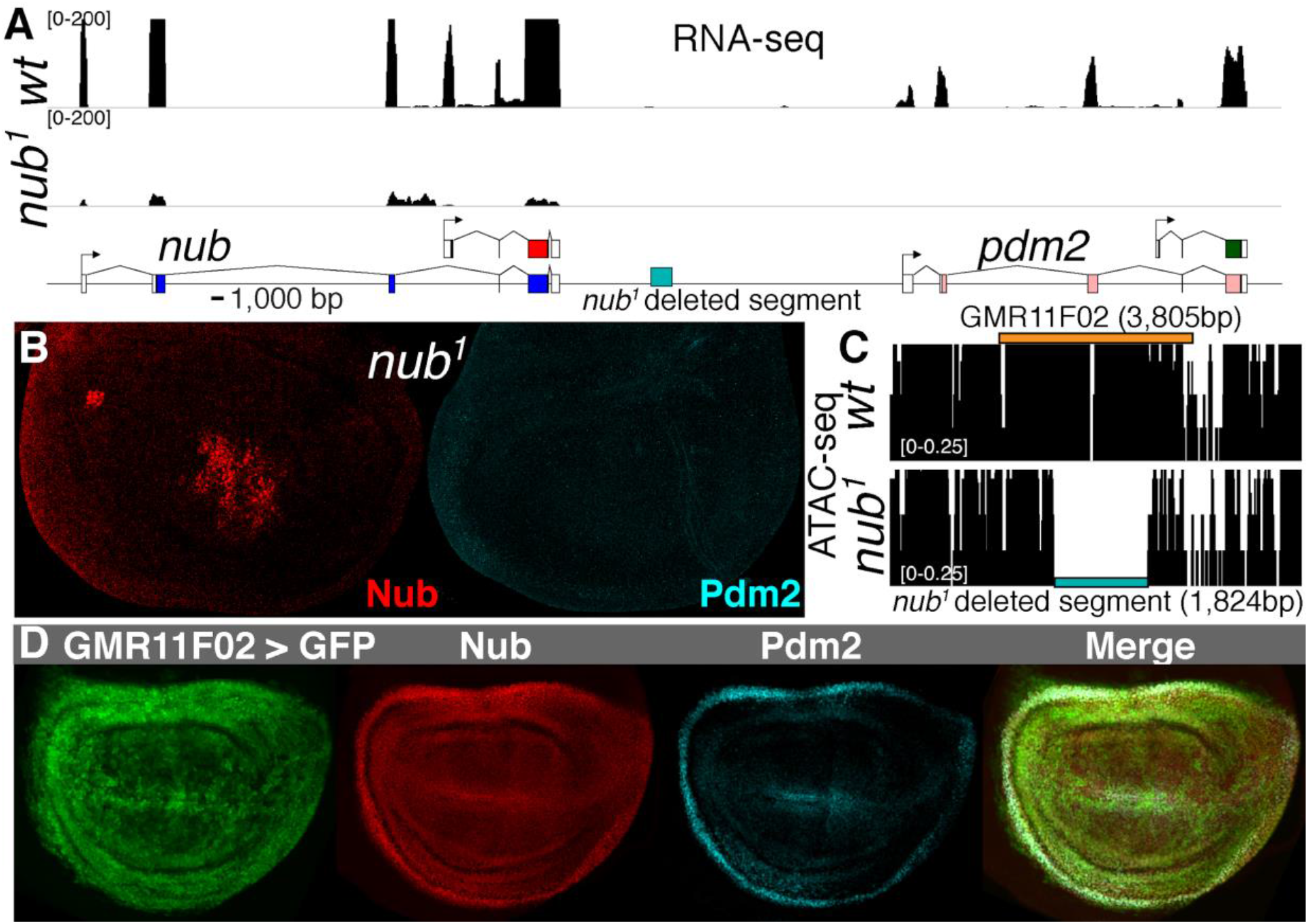
Additional characterization of the classic *nub*^*1*^ allele. (**A**) Transcriptomic analysis of the *nub*^*1*^ wing discs within the *nub*/*pdm2* genomic locus. Both *nub* and *pdm2* transcripts are reduced relative to the wild type (wt) *yw* strain. (**B**) Protein levels of Nub (left) and Pdm2 (right) in *nub*^*1*^ wing discs are reduced. Some staining of Nub is observed in the central pouch, as previously described, and consistent with the RNA-seq analysis. (**C**) ATAC-seq analysis revealed a lesion of DNA (blue box) in the intergenic region between *nub* and *pdm2* (see blue box in (**A**)). The location of reporter GMR11F02 overlapping this region is indicated (orange box). (**D**) Expression of a reporter construct containing GMR11F02 demonstrates that this segment is a wing domain enhancer overlapping the expression of Nub.

### *A wing enhancer element is deleted in the* nub^1^ *allele*

The *nub*^*1*^ allele has previously been shown to have an insertion of the 412 transposon into the promoter of one of two *nub* isoforms (Ng et al., 1995). However, this insertion is unlikely to explain the loss of expression of both paralogs. As part of our investigation of Nub function in the wing (discussed below) we performed Assay for Transposase Accessible Chromatin with high-throughput sequencing (ATAC-seq) in wild type and *nub*^*1*^ imaginal discs. ATAC-seq relies on Tn5 transposase insertion and cutting of genomic DNA, which occurs preferentially at open chromatin regions though also to a lower degree in all genomic regions (Buenrostro et al., 2013). Surprisingly, this analysis revealed a segment of DNA between *nub* and *pdm2* that had no ATAC-induced sequence reads from homozygous *nub*^*1*^ wing discs compared to wild type wing discs (Figure 3C). The most likely explanation is that this region is absent in the *nub*^*1*^ genome. To test this idea we directly sequenced the locus by Sanger sequencing, which confirmed that a sequence of 1,824 base pairs was missing in the *nub*^*1*^ genome (Figure S5). Thus *nub*^*1*^ contains a previously unknown 1.8 kb deletion in the intergenic region between *nub* and *pdm2*.

We next asked if the missing DNA fragment contains transcriptional regulatory activity. Previously, a large scale enhancer screen was performed and non-coding segments within the entirety of the *nub*/*pdm2* genomic locus were assayed for their ability to drive reporter gene expression (Jory et al., 2012). One fragment tested, GMR11F02, contains the entirety of the region deleted in the *nub*^*1*^ allele. When placed within a construct containing the *Drosophila* synthetic core promoter (DSCP) upstream of Gal4 coding sequence, UAS-driven reporter expression is observed throughout the developing wing appendage domain (Figure 3D). Thus a DNA fragment that includes the *nub*^*1*^ deletion has a previously unknown wing enhancer. Notably, among all cloned segments tiling the 140 kb *nub*/*pdm2* locus, GMR11F02 was the only fragment to drive an expression pattern similar to the *nub* expression domain (Jory et al., 2012). Below, the regulatory element contained within the GMR11F02 region will be referred to as the Nubbin-Pdm2-Wing (NPW) enhancer. In the next section we determine the extent to which NPW enhancer deletion contributes to the *nub*^*1*^ phenotype and its influence on expression of *nub* and *pdm2*.

### A nub/pdm2 shared enhancer is essential for wing development

The previous results show that the *nub*^*1*^ genotype is associated with a deletion of a wing enhancer contained within the GMR11F02 segment. As this allele also contains a 412 transposon insertion, and could have additional unknown abnormalities within the *nub*/*pdm2* locus, we next determined if the enhancer deletion was sufficient to cause the *nub*^*1*^ wing phenotype. To do this, we removed the NPW enhancer in an otherwise wild type background using CRISPR/Cas9. Two deletions were made: one corresponding closely to the missing segment in *nub*^*1*^ (NPW-small), and another that contains additional DNA present in the GMR11F02 segment (NPW-large) (Figure 4A-B). Flies containing the NPW-small deletion are homozygous viable with a strong wing phenotype similar to that of the *nub*^*1*^ animals (Figure 4C – compare with Figure 1E). The phenotype of adult flies containing NPW-large is indistinguishable from that of the smaller deletion (Figure 4C), suggesting that the region deleted in *nub*^*1*^ strain, corresponding to the NPW-small boundaries, removes the entire NPW enhancer. This result confirms that the primary cause of the *nub*^*1*^ phenotype is the absence of this enhancer element.

**Figure 4.**
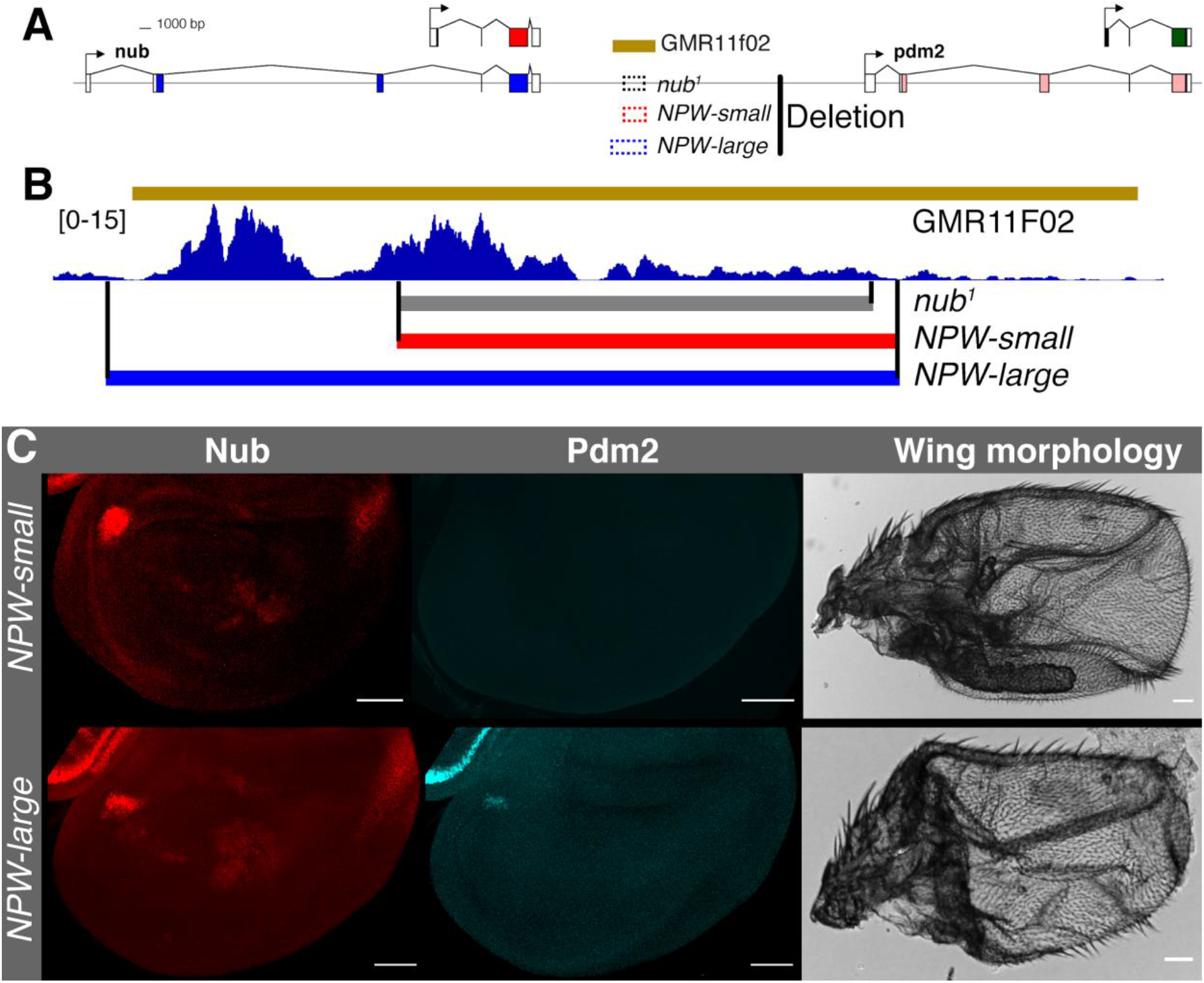
NPW is an essential wing enhancer shared by *nub* and *pdm2*. (**A**) Deletions corresponding to the lesion observed in the *nub*^*1*^ and CRISPR-mediated alleles (*NPW-small* and *NPW-large*) generated in this study within the *nub*/*pdm2* genomic locus are indicated. (**B**) Accessible chromatin in wild type wing discs in the genomic region overlapping GMR11F02. A large open chromatin peak is contained within the *nub*^*1*^ lesion and an additional adjacent peak is contained within the GMR11F02 region.The *NPW-small* deletion coordinates mimic the *nub*^*1*^ lesion while the NPW-large deletion removes both open chromatin regions. (**C**) *NPW-small* (top) and *NPW-large* deletion (bottom) phenocopies the *nub*^*1*^ allele both in regards to Nub and Pdm2 protein expression (left) and phenotype (right). Scale bar is 50 µm.

Because both *nub* and *pdm2* are sufficient for wing development, we next asked whether expression of both genes is dependent on this enhancer. Indeed, as in the *nub*^*1*^ background, both Nub and Pdm2 proteins are absent in the majority of the wing disc, with a small number of cells in the central region of the appendage domain retaining low levels of Nub (Figure 4C). Thus, this regulatory region is used by both paralogs during wing development. In distal hinge cells at the periphery of the wing progenitor domain the NPW enhancer is required for activation of both *nub* and *pdm2*. Under normal conditions in the rest of the appendage domain it preferentially activates *nub* due to repression of *pdm2*, but in the absence of *nub*, the NPW enhancer is also required to activate *pdm2*.

### Paralog specific response to NPW is due to repression by the Rotund transcription factor

Although Nub is essential for repression of *pdm2* in the central wing domain, additional spatial factors must be required as *pdm2* is not repressed in the *nub*+ cells at the periphery of the appendage domain. We looked for a potential candidate whose expression boundary in the wing disc is defined by where *pdm2* repression occurs. The transcription factor Rotund (Rn) is expressed in the central domain of the wing, encompassing the entire pouch and extending partially into the distal hinge (St Pierre et al., 2002). Co-staining a Rn-GFP allele tagged at the endogenous gene locus (Q. Li et al., 2015) and Pdm2 shows that the expression of Pdm2 and Rn is largely mutually exclusive (Figure 5A). To ask if Rn participates in *pdm2* repression, we examined Pdm2 levels in wing discs from flies containing a *rn* null allele. Consistent with a function for *rn* in repression of *pdm2*, we find de-repression of *pdm2* throughout the central wing disc in the absence of *rn* (Figure 5B). RNAi mediated knockdown of *rn* specifically in the posterior compartment driven by *engrailed* also causes derepression of Pdm2 specifically in posterior cells (Figure S6A). We next asked whether *rn* expression is dependent on *nub*, as this could potentially mediate the repression of *pdm2* by *nub*, and found that *rn* is expressed normally in the *nub*^*1*^ background (Figure S6B). Conversely, *nub* expression remains unaffected in the absence of *rn* (Figure 5B). Thus *nub* and *rn* do not regulate each other and are independently required for repression of *pdm2* in the central wing domain under normal conditions.

**Figure 5.**
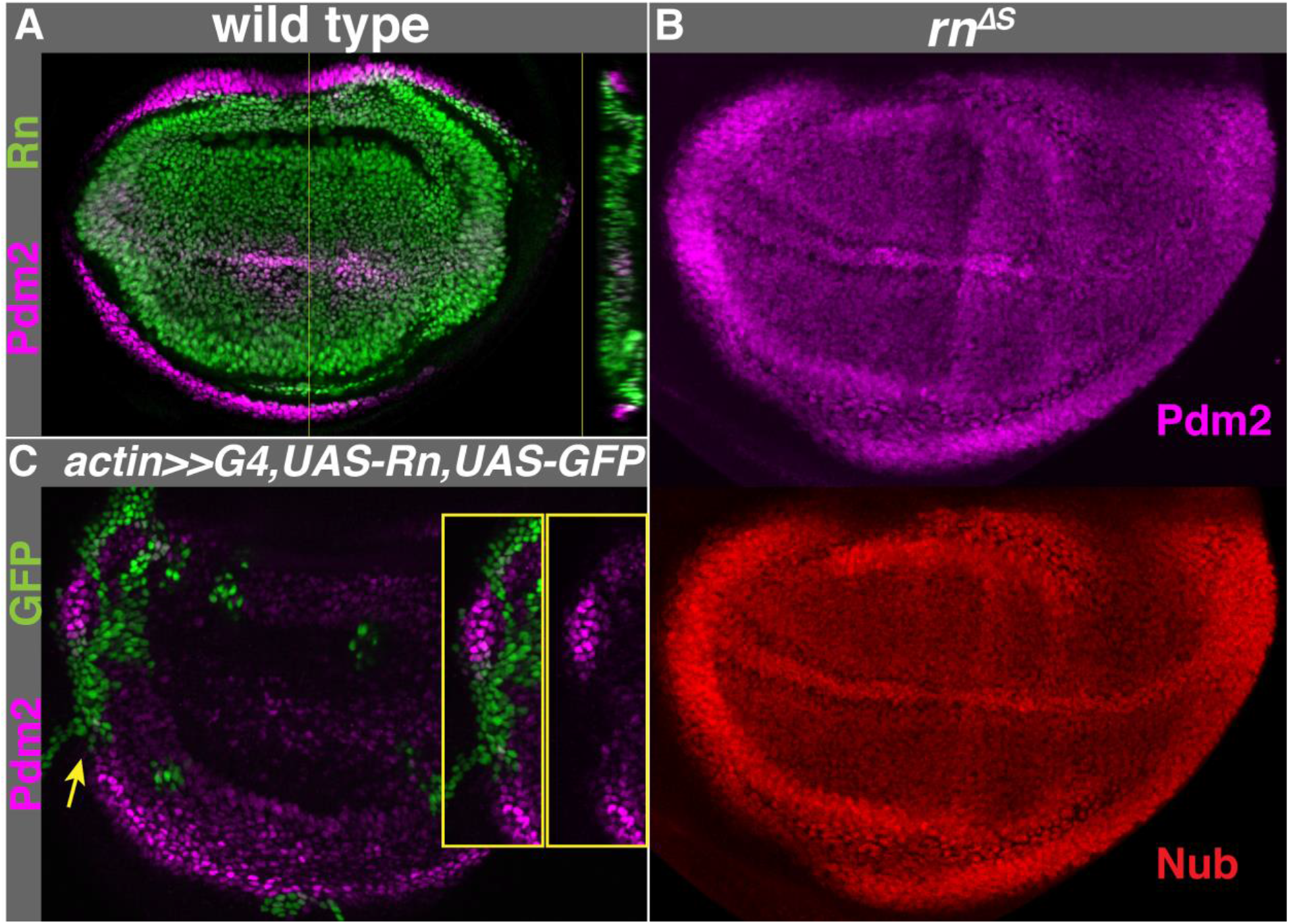
Repression of *pdm2* by Rotund. (A) *rotund*-GFP tagged allele (green) is expressed in the central wing domain adjacent to the Pdm2-high (magenta) wing periphery. Transverse section is shown (right). (**B**) Pdm2 (magenta) is derepressed in a *rn* null wing disc. (**C**) Heat shock induced clones ectopically expressing *rn* (visualized by GFP) cause repression of Pdm2 in the peripheral hinge domain.

If *rn* is the limiting factor that defines where *pdm2* is repressed, then ectopic expression of *rn* in the *pdm2*-expressing cells should be sufficient to repress *pdm2*. To test this we generated clones that ectopically express Rn. Expression of Rn in cells that normally express high levels of *pdm2* cause autonomous repression of *pdm2* (Figure 5C). In summary, *rn* is a *pdm2*-specific repressor that causes *nub* and *pdm2* to respond differently to the NPW enhancer.

### Rotund directly binds to the Pdm2 promoter

We next sought to identify the mechanism by which *rn* represses *pdm2*. As the NPW enhancer is shared by *nub* and *pdm2*, any repressive input into this regulatory element would likely affect both genes. Thus, we hypothesized that *rn* modifies *pdm2* expression through *cis*-acting regions outside of NPW or that *rn* represses *pdm2* indirectly. To ask if Rn binds to regions within the *nub*/*pdm2* genomic complex we performed Chromatin Immunoprecipitation with Sequencing (ChIP-seq) using the *rn-GFP* allele (Q. Li et al., 2015) with chromatin derived from otherwise wild type 3rd instar wing imaginal discs. Analysis of all regions of enriched ChIP-seq signal throughout the genome (i.e ChIP peaks) revealed a high degree of specific enrichment for the canonical Rn binding motif, suggesting that the experiment successfully identified Rn target sites (Figure S7A). Within the ∼140 kb *nub*/*pdm2* locus, there is a cluster of Rn peaks at the *pdm2* promoter, and weaker peaks at the promoters of *nub* and *ref2* (which overlaps the gene body of *nub* and is transcribed on the opposite strand) (Figure S7B). Given that our functional data show a specific effect of *rn* on *pdm2* expression, we hypothesized that the *pdm2*-specific response to NPW may be mediated by the cluster of Rn binding events within the *pdm2* promoter region (Figure 6A).

**Figure 6.**
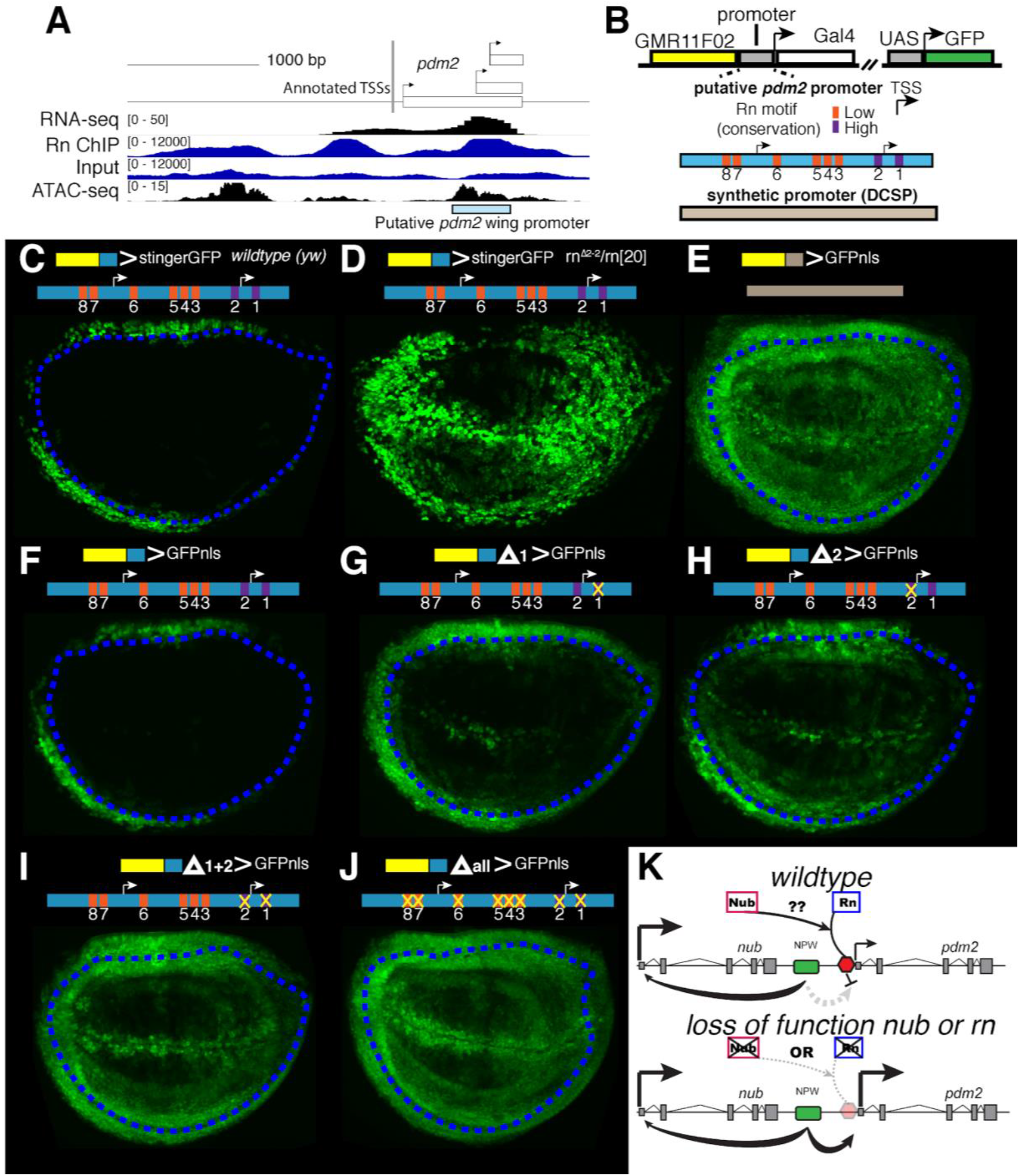
Rn directly represses *pdm2* through a promoter-localized silencer element. (**A**) *pdm2* promoter region with genome browser tracks of wing disc RNA-seq (top track), ChIP-seq enrichment of Rn and associated input control (second and third track, respectively) and ATAC-seq accessible chromatin (bottom track). Predicted wing promoter is indicated (blue box). (**B**) Diagram of enhancer-reporter constructs and different promoters used: putative *pdm2* promoter (blue) and Drosophila Synthetic Core Promoter (DSCP, tan). Consensus Rn binding motifs within the putative *pdm2* promoter shown and whether they are located in highly conserved sequence blocks is indicated by purple (yes) or orange (no). (**C-D**) Reporter expression from constructs containing GMR11F02 upstream of the *pdm2* promoter driving a UAS-GFPstinger(GFPnls variant) in wild type(**C**) and *rn* null(**D**) wing discs. Note that the peripheral hinge domain (where *pdm2* is normally expressed) is deleted in the *rn* null background. Therefore observed expression is likely all a result of derepression. (**E-J**) Reporter (UAS-GFPnls) expression of constructs containing GMR11F02 upstream of different promoter variants: (**E**) DSCP (**F**) wild type *pdm2* promoter (**G**) *pdm2* promoter: Rn site 1 mutant (**H**) *pdm2* promoter: Rn site 2 mutant (**I**) *pdm2* promoter: Rn site 1+2 mutant (**J**) *pdm2* promoter: All Rn sites mutant. Distal limit of Pdm2 protein expression is indicated by blue dashed line in all images with the exception of (**D**) where Pdm2 is uniformly expressed. (**K**) Model for paralog-specific suppression of NPW activity in wild type and mutant (*nub* or *rn*) conditions.

As Nub is also required for repression, we attempted a similar strategy to map its binding sites during wing development. To do so we made a *nub-GFP* knock-in allele using CRISPR/Cas9 and performed ChIP-seq on wing disc-derived chromatin. While this analysis revealed enrichment around the *pdm2* promoter, the global analysis failed to meet our criteria for a successful ChIP experiment. Specifically, the Nub motif enrichment in the genome-wide peak set was lower than that of general open-chromatin associated transcription factors such as GAF. Thus, whether Nub directly binds to the *pdm2* promoter remains an open question.

### The pdm2 promoter represses NPW activity

To test the hypothesis that the promoter region of *pdm2* is capable of modifying NPW enhancer activity, we utilized an enhancer-reporter system to test the effect of different promoters on NPW-dependent transcription. As discussed previously, GMR11F02, which contains NPW, drives reporter activity in the entire wing domain when placed upstream of a synthetic minimal promoter (Figure 3D and Figure 6E). When the synthetic promoter is replaced with a 444 nucleotide region comprising the putative *pdm2* promoter region, reporter activity recapitulates the pattern of the endogenous *pdm2* gene (Figure 6C). When observed in a *rn* null genetic background, in which *pdm2* expression is derepressed (see Figure 5B), reporter activity is also derepressed in most of the distal hinge and dorsal-ventral (DV) boundary, and to a lesser extent in the pouch cells surrounding the DV boundary (Figure 6D). This result is consistent with the ability of the *pdm2* promoter to modify the regulatory activity downstream of NPW in a Rn-dependent manner. We also observed the same response in the haltere imaginal disc, which is specified by an analogous genetic program as the wing, with an additional input by the *Hox* gene *Ultrabithorax* (*Ubx*) (Lewis, 1978; Loker et al., 2021) (Figure S8).

### *Repression of the* pdm2 *promoter by Rn is mediated by multiple binding sites*

To identify the Rn binding sites within the *pdm2* promoter that mediate repression we searched for matches to the Rn binding motif and assayed the effect of mutation of these sites on reporter activity. Because GMR11F02 has the capacity to function in all wing cells, mutation of repressive binding sites within the *pdm2* promoter should restore activity to the pattern observed when the synthetic promoter is used. Two Rn motifs proximal to the transcription start site are located within in a highly conserved block of 27 nucleotides conserved in all *Drosophila* species (referred to as motif #1 and 2, Figure S9). Furthermore, motif #1 and the surrounding 18 nucleotides are conserved within the distantly related housefly genome. An additional 6 motifs resembling Rn binding sites are present in the *D. melanogaster* genome that vary in their degree of conservation with other *Drosophila* species (Figure 6B and Figure S9). We generated four reporter variants to test the effect of these binding sites on NPW repression: 1) conserved motif #1 mutated, 2) conserved motif #2 mutated, 3) conserved motif #1+#2 mutated, and 4) all eight motifs mutated. Mutation of the motifs contained with the larger conservation block individually each results in expanded expression within the *rn*+ domain to varying degrees. Mutation of conserved motif #1 causes slight expansion of reporter activity distally in the hinge and derepression along the DV boundary (Figure 6G). In addition to increased activity within the *rn*+ domain, reporter expression is also stronger in cells outside of the *rn* expression domain, suggesting that this motif has additional Rn-independent functions (Figure 6G). Mutation of conserved motif #2 causes depression throughout the hinge domain, DV boundary, and some cells within the pouch (Figure 6H). When both site #1 and #2 are mutated together, a significant increase in reporter activity is observed throughout the *rn*+ domain (Figure 6I). The strongest derepression, as judged by the more uniform activity along the DV boundary and within the pouch, is observed when all eight motifs are mutated (Figure 6J). These results are consistent with an additive model, where individual motifs in the promoter region each contribute partially to repression of *pdm2* (Figure 6K). Even sites that are more poorly conserved contribute to the final level of repression.

### Repressive function of Nub/Pdm2 during wing development

The results presented thus far reveal how two paralogous genes have divergent expression patterns despite using a single, shared enhancer. To enhance our understanding of the selective pressures driving the divergent expression of these paralogs a better understanding of their function in wing development is required. Previous work has demonstrated a function for *nub* in antagonizing Notch signaling at the DV boundary by directly repressing several Notch-induced target genes (Neumann & Cohen, 1998). To systematically characterize how *nub* and *pdm2* function to influence wing appendage development on a genome-wide level we took two complementary approaches that both utilize ATAC-seq to identify the effect of Nub/Pdm2 on *cis*-regulatory accessibility. Enhancer activity is closely coupled to accessibility and quantitative differences in accessibility between related cell populations have been shown to be a reliable predictor of cell-specific enhancer activity (Janssens et al., 2022; Loker et al., 2021).

In the first approach we compared chromatin accessibility profiles using ATAC-seq datasets for two complementary domains in the wing imaginal disc: the *nub*-expressing wing progenitor cells and the *teashirt (tsh)*-expressing cells that will give rise to more proximal structures, the notum (body wall) and proximal hinge (Figure 7A) (Loker et al., 2021). Prior to specification of the wing progenitor domain as a result of *wg* and *dpp* signaling during late second instar, the entire imaginal disc expresses *tsh*. Transformation of distal (appendage) to proximal (notum) identity (or vise-versa) results from misexpression of proximal factors such as EGFR signaling or distal factors such as *wg* (Morata & Lawrence, 1975; Wang et al., 2000). Thus the differences in the chromatin and gene regulatory landscape in the appendage domain are likely driven by the loss of proximal-promoting factors and gain of appendage-promoting factors such as Nub/Pdm2 imposed upon a similar epithelial cell identity.

**Figure 7.**
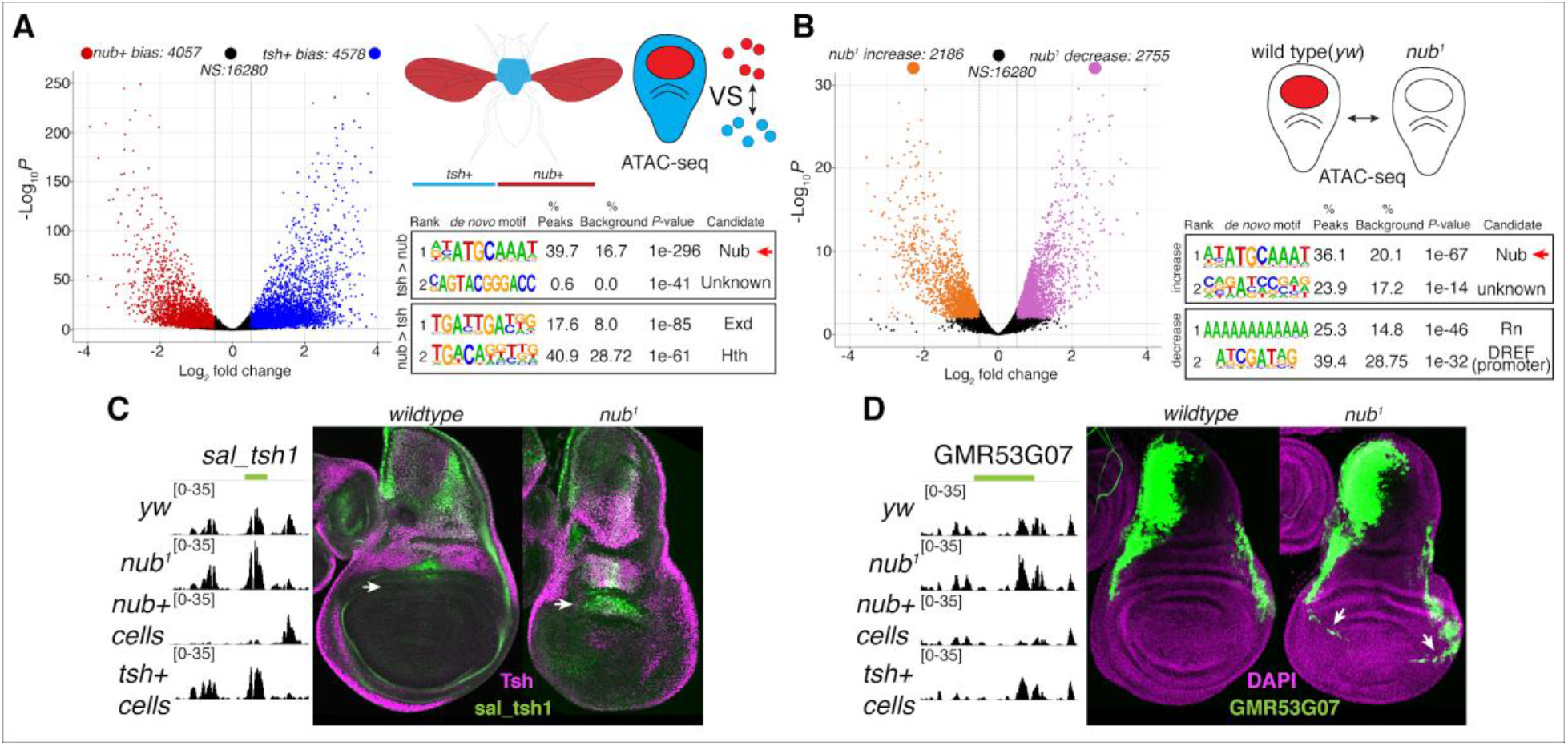
Influence of Nub/Pdm2 on wing chromatin landscape. (**A**) Comparison of ATAC-seq profiles from sorted *tsh+* and *nub+* wing disc cells. Right: Motif enrichment for regions with *nub+* bias or *tsh+* accessibility bias. (**B**) Comparison of ATAC-seq profiles of *yw* and *nub*^*1*^ wing discs. Right: Motif enrichment for regions that increase or decrease in *nub*^*1*^ discs. (**C**) Example regulatory region (called *sal_tsh1*) from the *spalt* genomic locus with *tsh* bias accessibility and increased accessibility in the *nub*^*1*^ background. Left: genomic ATAC tracks in indicated conditions. Right: Expression of reporter activity in wild type vs *nub*^*1*^ discs. (**D**) Example regulatory region (GMR73G07) from the *dlp* genomic locus with *tsh* bias accessibility and increased accessibility in the *nub*^*1*^ background. Left: genomic ATAC tracks in indicated conditions. Right: Expression of reporter activity in wild type vs *nub*^*1*^ discs.

This comparison revealed thousands of loci with increased or decreased accessibility in the *nub* domain relative to the *tsh* domain (Figure 7A). All previously described domain specific enhancers, such as the *vestigial* quadrant and *salm* pouch enhancers, have biased accessibility patterns, validating this approach for identifying region-specific regulatory elements (Guss et al., 2001; Kim et al., 1996; Loker et al., 2021). To distinguish differences directly associated with Nub/Pdm2 we performed separate motif searches within regions that have higher or lower accessibility in the *nub* domain. The most significantly enriched motif is a match to the consensus Nub/Pdm2 binding site, and this motif is specifically enriched in regions with less accessibility in the *nub* domain (Figure 7A). We also carried out the same analysis in sorted populations from the haltere imaginal discs, which yielded a similar enrichment of the same Nub/Pdm2 motif in regions with less accessibility in the *nub* domain relative to the *tsh* domain (Figure S10). The enrichment of the Nub/Pdm2 motif in genomic regions that have biased accessibility in the *tsh* domain suggests that Nub/Pdm2 functions to reduce accessibility in the *nub* domain, consistent with them functioning as repressors.

In the second approach we examined how chromatin accessibility changes in the absence of *nub*/*pdm2* by comparing the ATAC-seq profile of whole wing discs of wild type vs *nub*^*1*^ genotypes. Thousands of loci have an increase or decrease in accessibility, consistent with aberrant wing morphology upon loss of *nub*/*pdm2* (Figure 7B). To ask whether *nub*/*pdm2* is responsible directly for any of these differences we performed *de novo* motif searches in both categories. The most significantly enriched motif identified was a match to the consensus Nub/Pdm2 binding site (Figure 7B). Furthermore, enrichment of this motif is only present in regions that increase accessibility in the *nub*^*1*^ genotype. This observation supports a direct role for Nub/Pdm2 in decreasing chromatin accessibility. However, a confounding issue with these comparisons is that the *nub* domain is significantly smaller in the absence of *nub*/*pdm2*, which could contribute to the observed increase in the accessible regions in the *tsh* domain relative to the *nub* domain (Cifuentes & García-Bellido, 1997). Indeed, although *tsh*-biased peaks show an increase in accessibility in *nub*^*1*^ discs compared to wild type discs, *nub*-biased regions show a decrease in accessibility, and peaks without a *tsh*/*nub* bias are similar in *nub*^*1*^ and wild type conditions (Figure S11).

To ask if the changes in accessibility translates to differences in enhancer activity in the *tsh* and *nub* domains, we assayed enhancer reporter genes in wild type and *nub*^*1*^ wing discs. We characterized two enhancers with a consensus Nub/Pdm2 motif that show *tsh*-biased accessibility in wild type tissue and increased accessibility in the *nub*^*1*^ background. These enhancers (*sal_tsh1* and GMR53G07 (Jory et al., 2012)) both displayed a *tsh*-biased expression pattern in wild type imaginal discs and derepression within the appendage domain in *nub*^*1*^ wing discs, consistent with Nub/Pdm2 functioning as a repressor of both enhancers (Figure 7C-D). We further characterized the response of *sal*, the gene presumably regulated by *sal_tsh1*. Antibody labeling revealed derepression of Spalt in the distal hinge region of *nub*^*1*^ wing discs coinciding with derepression of the *sal_tsh1* activity (Figure S12A-B). Additionally, ectopic expression of Nub in the *tsh* domain represses *sal*, supporting the idea that Nub/Pdm2 represses the activity of enhancers that are normally active in the *tsh* domain (Figure S12C).

## Discussion

In this paper we described a single essential wing enhancer element that is interpreted in two different ways by two paralogous genes, *nubbin* and *pdm2*. This difference results in *nub* being the predominant paralog used during wing development under wild type conditions, while *pdm2* is suppressed by *nub* and *rn*. Upon loss of *nub*, upregulation of *pdm2* permits functional compensation and normal wing development, revealing a redundant function for *pdm2* and *nub* in wing development. As such this system serves as a backup to ensure robust wing development, while keeping the combined Nub/Pdm2 protein levels at the correct level during normal conditions. A function for gene duplicates in providing robustness to genetic and environmental perturbations has been demonstrated in numerous circumstances, but the mechanism by which compensation is achieved is often unknown (Gu et al., 2003; Osterwalder et al., 2018).

### Evolution of gene regulation in the context of shared enhancers

One potential explanation for the shared expression between duplicated genes, particularly tandem duplicates, is the co-dependence on shared regulatory elements (Lan & Pritchard, 2016; Quintero-Cadena & Sternberg, 2016; Williams & Bowles, 2004). Shared regulatory elements have long been implicated in the nested expression patterns of *HOX* paralogs (Gould et al., 1997). More recently shared enhancers have been implicated in a wider array of paralogous genes as a result of large-scale reporter screens and chromosome conformation capture (3C) approaches, which identified enhancers that putatively regulate the promoters of many co-expressed paralogs (Ghavi-Helm et al., 2014; Jory et al., 2012; Kvon et al., 2014; Lan & Pritchard, 2016; Symmons et al., 2014). While these techniques suggest the widespread utilization of shared enhancers, in most cases functional validation is lacking and the existence of multiple enhancers that separately control both genes cannot be ruled out. Additionally, although there are several models to account for subsequent evolutionary steps following gene and enhancer duplication, it is less clear whether and how context-specific divergence in expression can occur when paralogs are under the control of shared enhancers. Interestingly, both enhancer reporter screens and 3C assays used in the *Drosophila* embryo have suggested the existence of many shared enhancers within the *nub*/*pdm2* that function in both the ectoderm and central nervous system (CNS) (Ghavi-Helm et al., 2014; Kvon et al., 2014; Ross et al., 2015). Thus there may be many shared regulatory elements, in addition to the wing NPW element described here, used by *nub* and *pdm2* throughout development.

Our results provide an interesting comparison with previous work on *nub*/*pdm2* function in the CNS. Although the enhancer(s) governing their expression is not known, *nub* and *pdm2* are co-expressed in the RP2 motor neurons of the embryonic CNS (Yeo et al., 1995). Loss of either paralog results in a reduction in the number of RP2 neurons, though *pdm2* has a more pronounced effect, while removal of both genes causes the complete loss of these neurons (Yeo et al., 1995). Thus in this case the expression of both paralogs is required for complete specification of RP2 neurons, consistent with a dosage subfunctionalization model in which expression of both paralogs has been reduced relative to the ancestral state (Gout & Lynch, 2015). In contrast to these observations, here we show that in the majority of wing progenitor cells *pdm2* is actively repressed such that in normal conditions only *nub* is present. The presence of a promoter-localized silencer element requiring *rn* and, via direct or indirect input, *nub*, permits the repression of *pdm2* by *nub*. Importantly, this repression is tissue-specific because it also requires the wing TF, Rn. As a consequence, *nub* does not repress *pdm2* through this silencer outside of the wing, allowing co-expression in other contexts such as the wing hinge and embryonic CNS. Furthermore, the dependence on *nub* for the silencer to function allows *pdm2* to respond to the NPW enhancer when *nub* levels are compromised. Thus, multiple mechanisms can evolve in distinct cell types leading to expression divergence of tandemly duplicate genes.

### Flexibility of gene regulation and evolution of silencer elements

The results described here prompt speculation as to how the divergent patterns of *nub*/*pdm2* in the wing may have evolved. Given that *nub*/*pdm2* is a marker for wing development in all winged insects, including species that diverged prior to duplication, it is likely that uniform expression similar to the *nub* pattern is the ancestral state. Thus repression of *pdm2*, through the promoter-proximal silencer element identified here, likely represents an evolutionary novelty that arose at some point post-duplication. The high degree of conservation for two Rn binding sites in the *pdm2* silencer element suggests this divergence may have been fixed shortly after the ancestral gene was duplicated. The acquisition of this silencer permitted conditional separation of *nub*/*pdm2* transcripts while maintaining the ability of *pdm2* to function in a compensatory manner when needed. Alternatively, if duplicated enhancers were used to independently regulate *nub* and *pdm2, rn*/*nub* repression of *pdm2* could have been mediated via a *pdm2*-specific enhancer. However, the dependence on a single enhancer in principle necessitates repression occurring independently of the shared enhancer. One mechanism that has been described to mediate promoter selectivity is an inherent preference of enhancers for specific promoter motifs, such as TATA or DPE motifs (Juven-Gershon et al., 2008; Ling et al., 2019). However if such a mechanism was operating for *nub*/*pdm2*, it would be inconsistent with our observation that NPW activates both promoters in part of the distal hinge, and the ability of *pdm2* to respond to the NPW enhancer in the absence of *nub*. The mechanism identified here for conditional repression of *pdm2* thus provides an elegant solution that may be widely applicable to other cases of shared enhancers, or gene regulation in general.

Notably, such promoter-proximal silencers that function to modify distal enhancers have not been identified previously, though transgene experiments have established that repression of enhancers can act over long distances (Cai et al., 1996). In contrast, long-range silencer elements have been implicated in previous studies. For example, the analysis of transcriptional regulation of *ebony*, which encodes an enzyme that contributes to yellow cuticle in the developing epidermis of *Drosophila* (Rebeiz et al., 2009), showed that an upstream enhancer collaborates with a silencer element in the *ebony* intron. An additional silencer is required for sex-specific repression of *ebony* and pigmentation patterns. The repeated evolution of this silencer element through accumulation of inactivating and spatial pattern-affecting mutations in different *Drosophila* species has contributed to species-specific pigmentation patterns (Johnson et al., 2015). Our analysis suggests that similar acquisition of silencer activity allowed the expression of *pdm2* to diverge from *nub* in the *Drosophila* species that depend on the shared NPW enhancer. Notably, the additive nature of Rn binding sites contributes to spatial differences in repression. We speculate that this is a result of spatial differences in the activating potential of the NPW wing enhancer. In regions where activity is stronger, more repressive input is required, whereas cells where the enhancer is weaker require less repressive input. In further support of this idea, when GMR11F02 is upstream of a synthetic promoter, although it is active throughout the wing domain, its activity is strongest in the distal hinge and DV boundary cells, the same cells that are most sensitive to derepression of the *pdm2* promoter when individual Rn binding sites are mutated. Thus spatial control of silencer activity may gradually evolve through the additive accumulation of repressor binding sites, and the amount of repression will be inversely proportional to the strength of the enhancer.

### The role of Nub/Pdm2 during wing development

Our results suggest that Nub/Pdm2 proteins function in the wing disc as repressors of transcriptional enhancers by decreasing chromatin accessibility. Interestingly, Nub has been previously shown to antagonize several genes downstream of the Notch signaling pathway (Neumann & Cohen, 1998). We propose that Nub/Pdm2 has a broader role as a repressor functioning to reduce the activity of many *cis*-regulatory enhancers within the appendage domain relative to the proximal domain that gives rise to the body wall. As the appendage and body wall progenitors share a common origin and rely on shared signalling molecules such as Dpp and Notch, *cis*-regulatory targets downstream of these pathways may be predisposed to respond equally in both domains. As such, an appendage-specific transcription factor may be required to ensure that enhancers specifically required in the proximal domain are not activated in the appendage domain. As Nub is the only known transcription factor expressed in all wing appendage cells (comprising the distal hinge and wing blade) throughout development, it is well suited to perform such a role.

## Methods

### Reporter constructs

The GMR11F02 reporter construct was generated previously by inserting the genomic coordinates:chr2L:12,634,617-12,638,421 (dm6 assembly) into the pBPGUw vector containing the Drosophila synthetic core promoter (DSCP) upstream of the Gal4 coding sequence. This construct was re-injected into the attP40 landing site (this study) by Rainbow Transgenic injection service. To exchange the DSCP with the endogenous *pdm2* promoter, the sequence was synthesized either corresponding to the reference genome or with indicated *rn* motif mutants by Genewiz containing restriction sites for FseI and KpnI (see Key Resource table). The DSCP was removed from GMR11F02 using the FseI and KpnI restriction sites, and replaced with the *pdm2* promoter. All plasmids were inserted into the attp40 landing site for this study using standard phiC31 injections performed by Rainbow Transgenic Flies injection service

### Generation of NPW enhancer deletions

Preparation of materials for Cas9-mediated deletions were performed as previously described(Alexandre et al., 2014). Potential gRNA target sites (two sites per deletion) were predicted using the flyCRISPR bioinformatic tool and subsequently sanger sequencing on the genomic region surrounding target sites was performed to confirm their presence in the Cas9-expressing stock (Rainbow Transgenic Flies: line#55821). gRNAs (see Key Resources table) were cloned into the pCFD5 vector. For donor constructs the TVattp-Pax-Cherry was used and homology arms were inserted using standard restriction enzyme mediated cloning with primers listed (see key resource table). For the NPW small and large deletion the same gRNA and Homology arm on one side was used. Injection was carried out by Rainbow Transgenic using the Cas9-expressing fly strain. Screening for successful deletion was performed using Pax-Cherry expression, and subsequently gDNA was extracted and sanger sequencing performed to confirm boundaries of Cas9-mediated lesions. Stocks were established from lines containing the correct targeting and subsequently flies were crossed to a Cre-expression strain to remove Pax-Cherry marker using flanking LoxP sites contained in the targeting vector. The resulting stocks contain a single attP and LoxP site in place of the deleted fragment.

### Immunohistochemistry

Immunohistochemistry was performed using standard practices as previously described(Loker et al., 2021).

### Ectopic expression of Rotund

To make Rn-expressing clones female flies from a *yw hs-flp*^*1*.*22*^*;UAS-FRT*.*STOP*.*FRT-Gal4* strain were crossed to males containing *UAS-Rn* (Bloomington:7403). 72 hours after egg laying, larvae from this cross were placed at 37C for 10 minutes to induce clones and analyzed 48 hours later by antibody staining of late 3rd instar.

### ChIP-seq

ChIP-seq was performed as described (Loker et al., 2021) previously on 3^rd^ instar larvae containing a GFP knock-in of the endogenous *rn* coding sequence(Q. Li et al., 2015). Briefly, 3^rd^ instar larval heads were dissected and inverted in PBS on ice. Heads were fixed for 20 minutes in 1.8% PFA in crosslinking medium (10 mM HEPES, pH=8.0; 100 mM NaCl; 1 mM EDTA, pH=8.0; 0.5 mM EGTA, pH=8.0) at room-temperature with rotation, and subsequently quenched (Quench solution: 1xPBS; 125 mM glycine; 0.1% Triton X-100). Fixed-heads were then washed 2X in buffer A (10 mM HEPES, pH=8.0; 10 mM EDTA, pH=8.0; 0.5 mM EGTA, pH=8.0, 0.25 % Triton X-100) and 2X buffer B (10 mM HEPES, pH=8.0; 200 mM NaCl; 1 mM EDTA, pH=8.0; 0.5 mM EGTA, pH=8.0; 0.01% Triton X-100) 10 minutes each at 4°C. Wing discs were then dissected and placed in sonication buffer (10 mM HEPES, pH = 8.0 ;1 mM EDTA, pH = 8.0; 0.5 mM EGTA, pH = 8.0, 0.1 % SDS). Chromatin sonication was performed using a Covaris S2 instrument at settings (105W; 2 % Duty; 15 minutes).

Sonicated chromatin was brought to 1X mild-RIPA (10 mM Tris-HCl, pH=8.0; 1 mM EDTA, pH=8.0; 150 mM NaCl; 1% Triton X-100) concentration and pre-cleared with Dynabeads for 1 hour at 4 °C with rotation. Pre-clearing beads were removed and antibody was added for incubation overnight. Dynabeads were added and incubated for 3 hrs. Bead bound antibody-chromatin complexes were washed as follows 2X RIPA LS (10 mM Tris-HCl, pH=8.0; 1 mM EDTA, pH=8.0; 150 mM NaCl; 1% Triton X-100; 0.1 % SDS; 0.1 % DOC), 2X RIPA HS (10 mM Tris-HCl, pH=8.0; 1 mM EDTA, pH=8.0; 500 mM NaCl; 1% Triton X-100; 0.1 % SDS; 0.1 % DOC), 1X LiCl (10mM Tris-HCl, pH=8.0; 1 mM EDTA, pH=8.0; 250 mM LiCl; 0.5 % IGEPAL CA-630; 0.5 % DOC), 1X TE (10 mM Tris-HCl, pH=8.0; 1 mM EDTA, pH=8.0). Samples were then treated with RNAse and proteinase K, and chromatin was isolated using phenol-chloroform.

An anti-GFP (ab290, Abcam; 1:300 dilution for IP) was used to perform chromatin precipitation. ChIP-seq libraries were made following the NEBnext UltraII kit (NEB) and associated protocol. Libraries were sequenced using a 75-cycle high output with single end sequencing using an Illumina Nextseq. Two replicates were performed.

### RNA-seq

For *nub*^*1*^ and wild type control samples, RNA was extracted from dissected whole discs using TRIzol and purified using Zymo Direct-zol RNA Microprep kit. Ten discs were used for each replicate. Libraries were prepared using NEBnext Ultra II Directional RNA-seq kit and sequenced using a 75-cycle single-end high output run with an Illumina Nextseq. Three replicates were performed for each experiment.

### ATAC-seq

For *nub*^*1*^ and wild type control samples, whole wing discs were dissected and subjected to the ATAC-seq protocol as previously described (Buenrostro et al., 2013; Jacobs et al., 2018). Three discs were used for each replicate. Libraries were sequenced using a 75-cycle high output with paired end sequencing using an Illumina Nextseq. Two replicates were performed for all ATAC-seq experiments.

### RNA-seq data processing

Reads were mapped using HISAT2 to the dm6 genome assembly. Mapped reads were then filtered for map quality using SAMtools (H. Li et al., 2009). Genome-track files were created using Deeptools (BamCoverage; RPGC normalization) (Ramírez et al., 2016).

### ATAC-seq data processing

Reads were mapped using Bowtie2 to the dm6 genome assembly. Mapped reads were then filtered for map quality using SAMtools (H. Li et al., 2009) and duplicates (Picard MarkDuplicates). Genome-track files were created using Deeptools (BamCoverage; RPGC normalization) (Ramírez et al., 2016).

Differential analysis was performed using DESeq2(Love et al., 2014) on a common interval of 24,915 peaks generated by merging ATAC-seq peaks called by MACS2 (callpeak --nomodel -- call-summits)(Zhang et al., 2008). Cut off used for calling differential accessibility was Log_2_Fold change > 0.5 and adjusted p-value (padj) < 0.05.

### ChIP-seq data processing

Reads were mapped using Bowtie2 to the dm6 genome assembly. Mapped reads were then filtered for map quality using SAMtools(H. Li et al., 2009) and duplicates (Picard MarkDuplicates). Peaks were called using MACS2 callpeak function with default parameters(Zhang et al., 2008). Genome-track files were created using Deeptools (BamCoverage; RPKM normalization)(Ramírez et al., 2016).

### Motif analysis

*De novo* motifs were discovered using Homer (findmotifsgenome.pl) (Heinz et al., 2010). For ATAC-seq data the entire peak was used to search for enriched motifs (option: -size given) and all ATAC peaks (minus the queried group) were used to calculate background enrichment. For ChIP-seq a default 200bp window around the peak center was used.

### Somatic CRISPR

To generate wing discs with somatic *nub* knockout, female flies containing *UAS-Cas9*.*M*(Port et al., 2020);*hdc*.*G4* were crossed to flies containing *UAS-nub*.*gRNA* (containing two gRNAs within the pCFD6 plasmid targeting coding exons shared in all *nub* isoforms, Vienna Drosophila Resource Center: 341604). When performed in *pdm2* null background: *UAS-Cas9*.*M,pdm2*^*E46*^*;hdc*.*G4* and *pdm2*^*E46*^,*UAS-nub*.*gRNA* stocks were used.

## Supporting information

supplemental_oligos

## Acknowledgements

We would like to thank all members of the Mann lab and Peter Andolfatto for helpful discussions during the course of this project and to David Stern for comments on the manuscript; the Bloomington Drosophila Stock Center for reagents; Flybase for information related to alleles used here; Pelin Volkan for the *rn*-GFP stock; Wes Grueber for fly reagents; Cyrille Alexandre and J.P Vincent for CRISPR targeting reagents; Fillip Port for CRISPR fly reagents. This work was supported by NIH grant R35GM118336 awarded to R.S.M.

## Author contributions

Conceptualization, R.L. and R.S.M.; Methodology, R.L. and R.S.M.; Investigation, R.L.; Formal Analysis, R.L.; Visualization, R.L.; Writing – Original Draft, R.L. and R.S.M.; Writing – Review & Editing, R.L. and R.S.M.; Funding Acquisition, R.S.M.

## Data availability

All imaging data and fly stocks associated with this study are available upon request. Genomic data generated previously corresponding to sorted wing and haltere imaginal disc populations is available within the GEO database with the accession **GSE166714**. Sequencing data generated from this study using RNA-seq(yw and nub^1^) and ATAC-seq(yw and nub^1^) will be made available on GEO upon publication.

## Notes

### Competing Interest Statement

The authors have declared no competing interest.

