## supplemental_figures for "A promoter-proximal silencer modifies the activity of a shared enhancer to mediate divergent expression of *nub* and *pdm2* paralogs in wing development"

### Supplementary figures

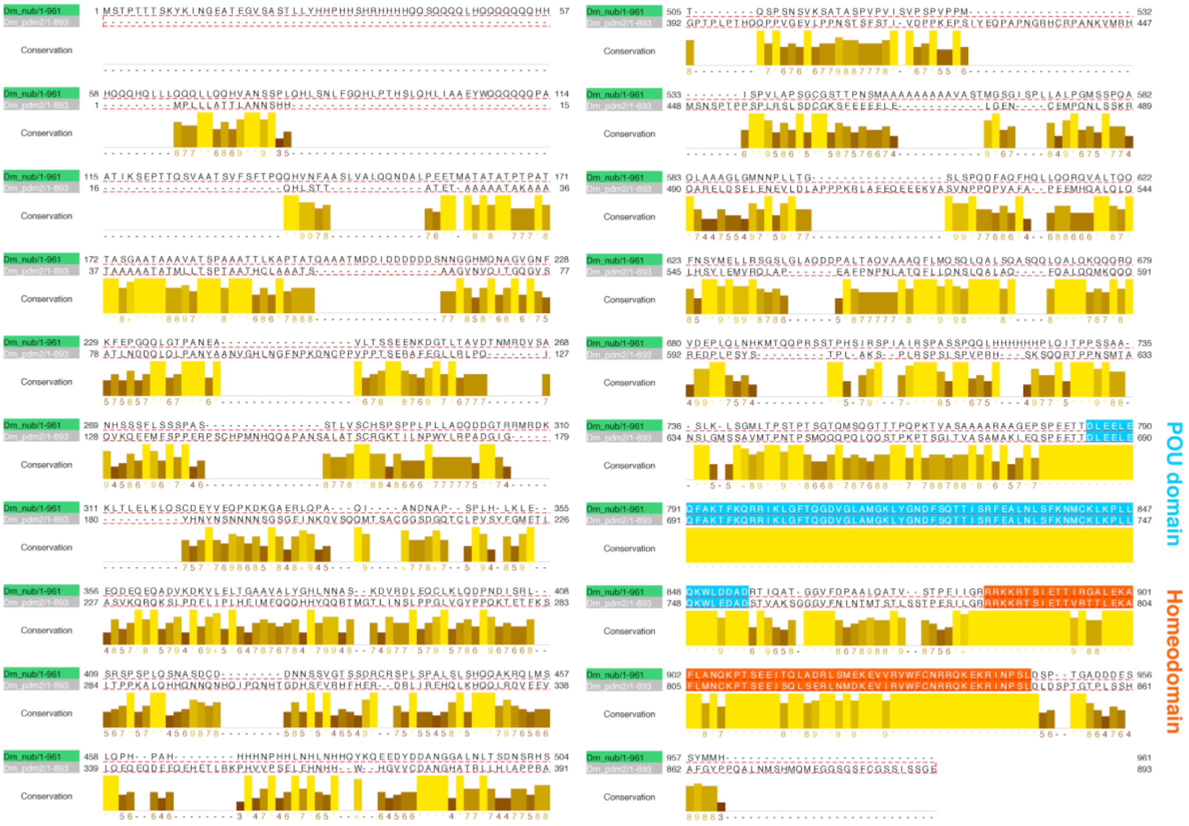


**Figure S1**

Protein alignment of long isoform of Nub (Top) and Pdm2 (Bottom). Highly conserved domains corresponding to POU-domain (blue) and Homeodomain (orange) are indicated.

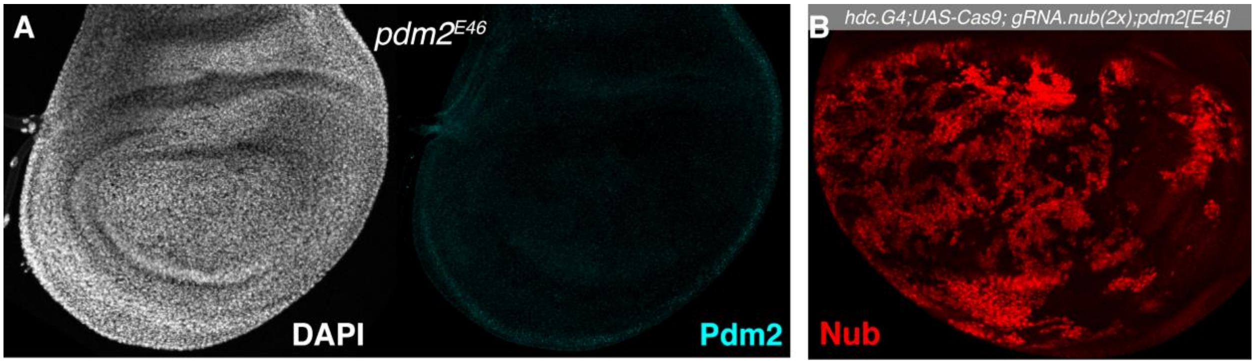


**Figure S2**

(A) *pdm2*$^{E46}$ wing disc stained with anti-Pdm2

(B) Targeting *nub* with somatic CRISPR results in mosaic tissue with patches of *nub*+ and *nub*- cells

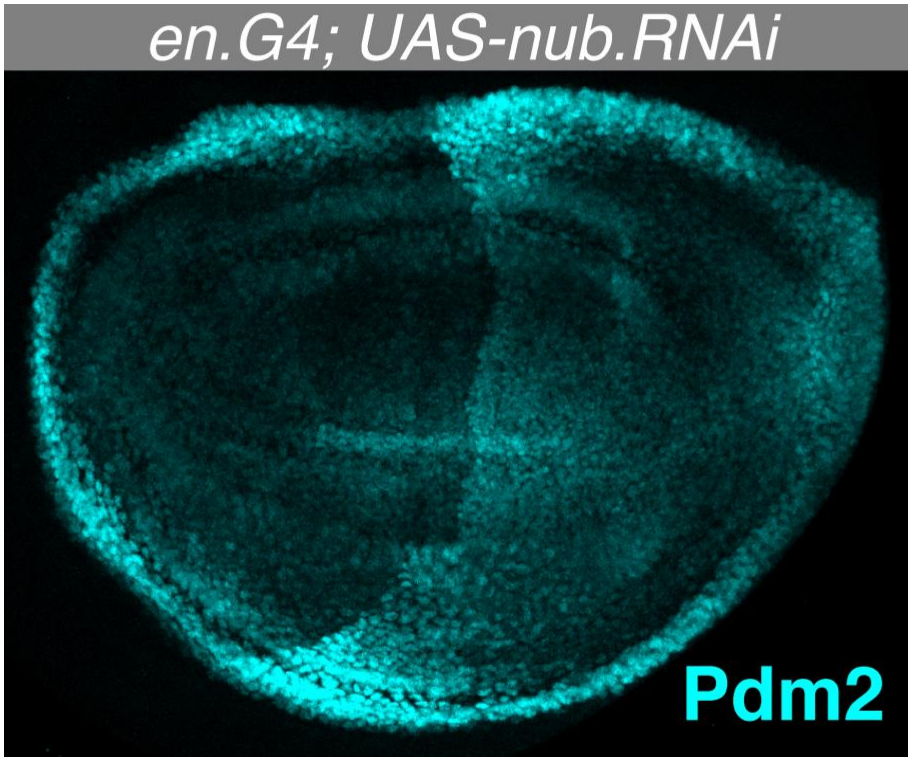


**Figure S3**
RNAi-mediated knockdown of *nub* using the *en.G4* driver in the posterior compartment results in posterior-specific upregulation of Pdm2 protein.

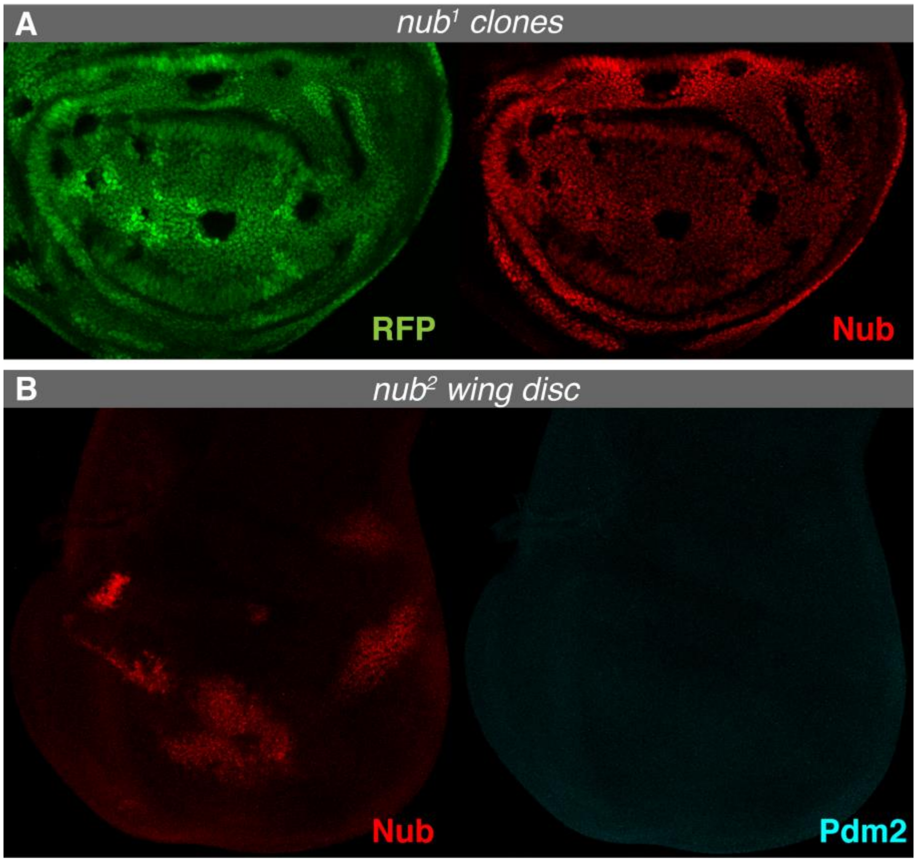


**Figure S4**

(A) Clonal analysis of the *nub*$^{1}$ allele in which Nub protein levels (Red) of wildtype (GFP+) cells can be compared with homozygous mutant (GFP-).

(B) Nub and Pdm2 protein levels are reduced in homozygous *nub*$^{2}$ wing discs.

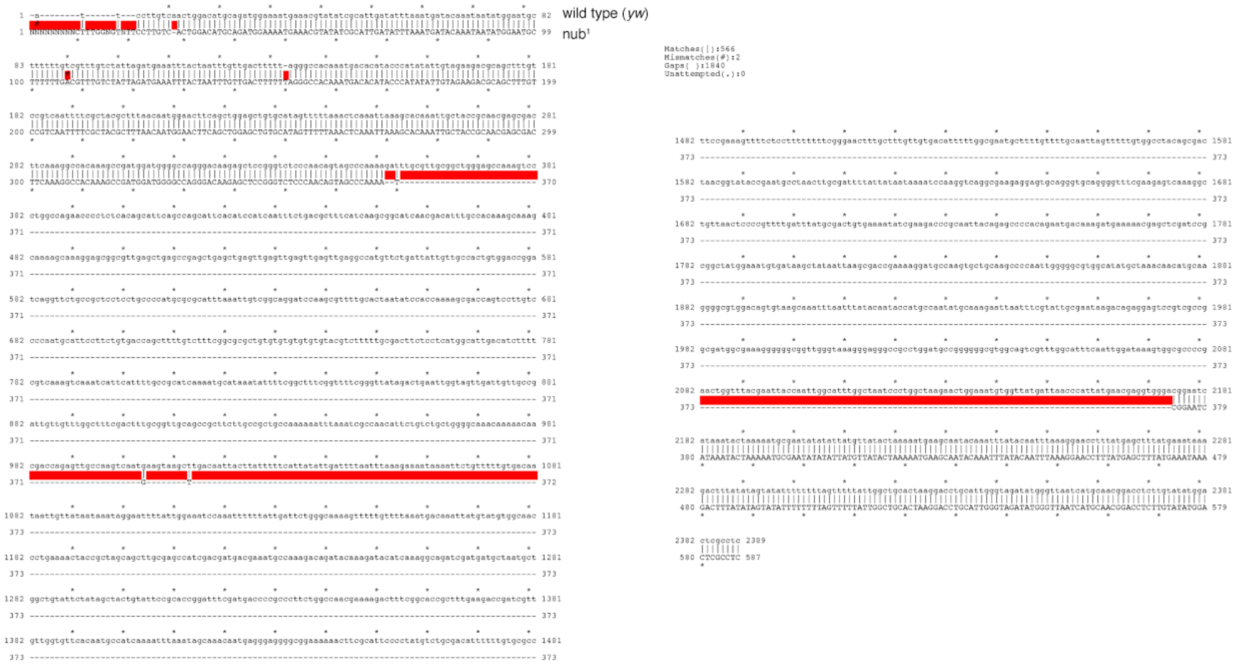


**Figure S5**

Alignment of Sanger sequencing of genomic DNA from wild type and homozygous *nub*[1] animals within the region surrounding the deleted segment.

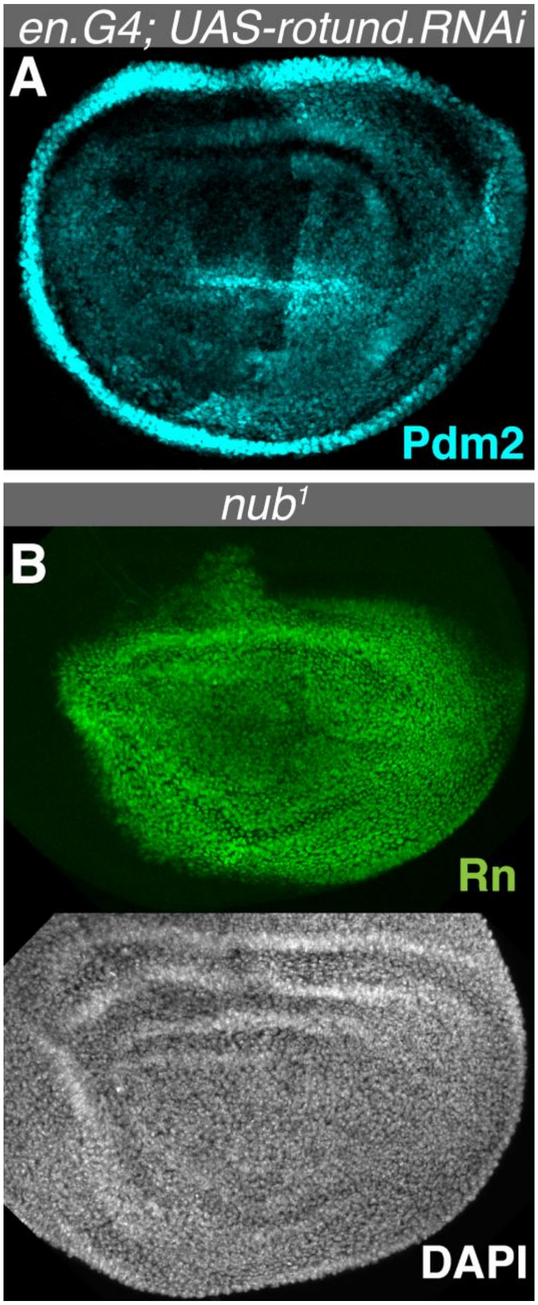


**Figure S6**

(A) RNAi-mediated knockdown of *rn* using the *en.G4* driver in the posterior compartment results in posterior-specific upregulation of Pdm2 protein.

(B) Expression of the *rn*-GFP allele (Top) in homozygous *nub*$^{1}$ wing discs. DAPI channel image is also shown (bottom)

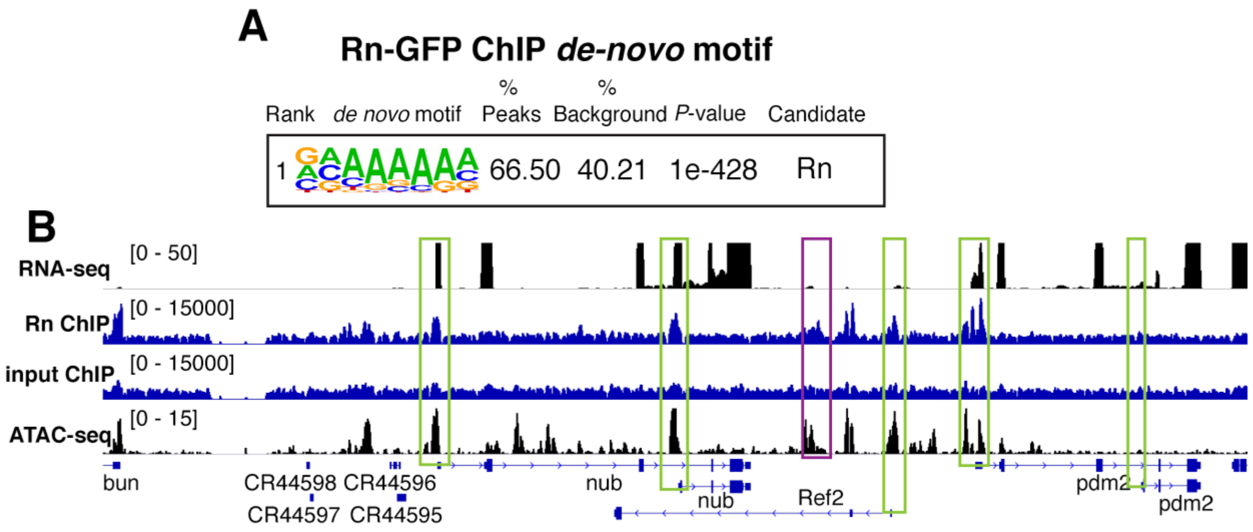


**Figure S7**

(A) Genomic tracks showing indicated experiments in the *nub*/*pdm2* genomic locus. Colored boxes indicate promoter regions (green) and the GMR11F02 region (purple).

(B) *de-novo* motif enrichment within Rn-GFP ChIP peaks from wing imaginal discs.

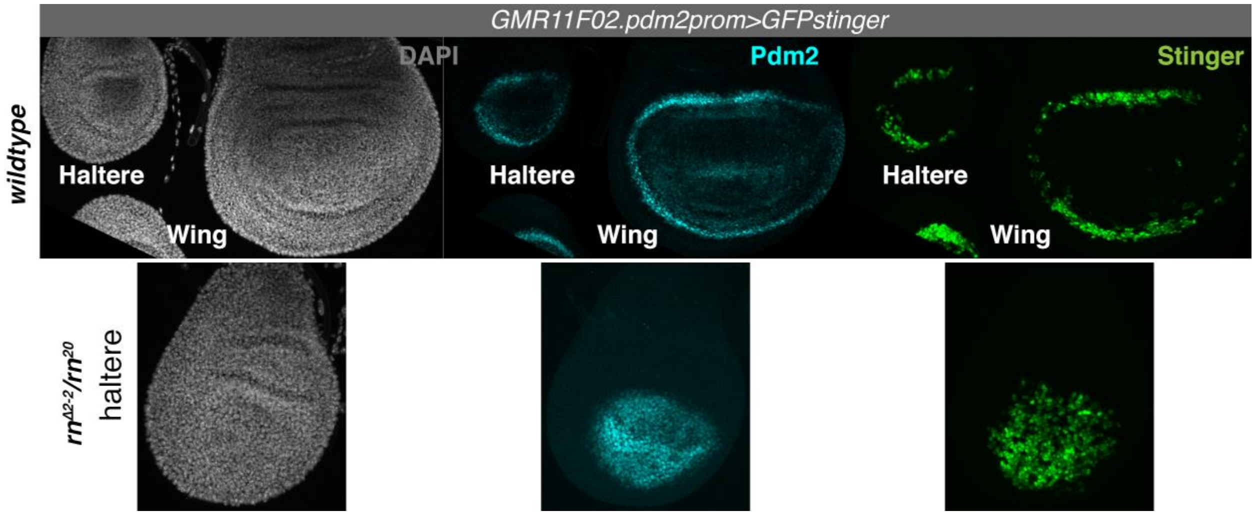


**Figure S8**

Activity of GMR11F02 is modified by the *pdm2* promoter similarly in wing and haltere imaginal disc. Top: Pdm2 protein levels (cyan) and expression of reporter activity (UAS-GFPstinger) driven by R11F02 upstream of *pdm2* promoter in the wing and haltere. Bottom: Pdm2 protein levels and reporter activity in *rn* null haltere discs. Derepression of Pdm2 and reporter activity is observed similar to that of the wing disc (see Figure 5B and Figure 6D).

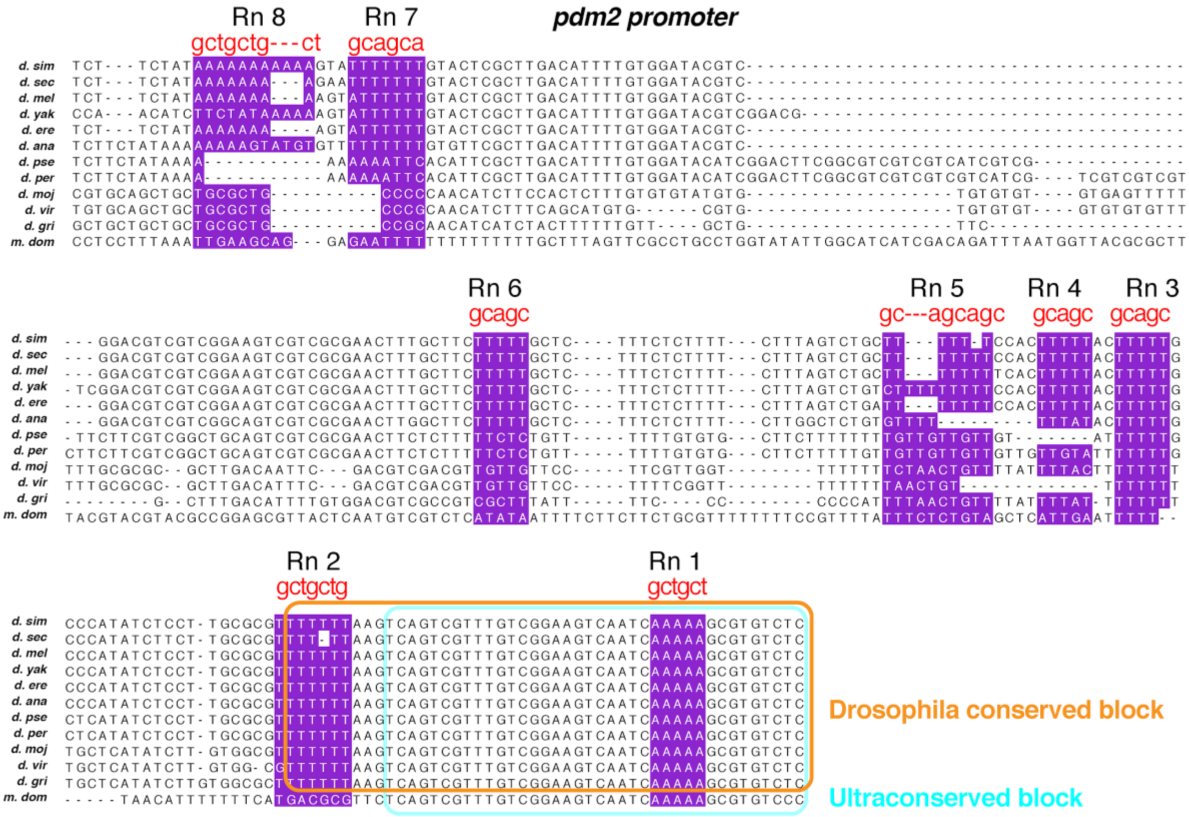


**Figure S9**

Conservation of rn binding sites(purple). Sequence mutations used are shown in red above alignment. A block of conserved sequence (orange) containing sites #1 and 2 is shared in *Drosophila* species, and part of this segment (blue), containing site #1, is conserved in the housefly (*m. domestica*) genome.

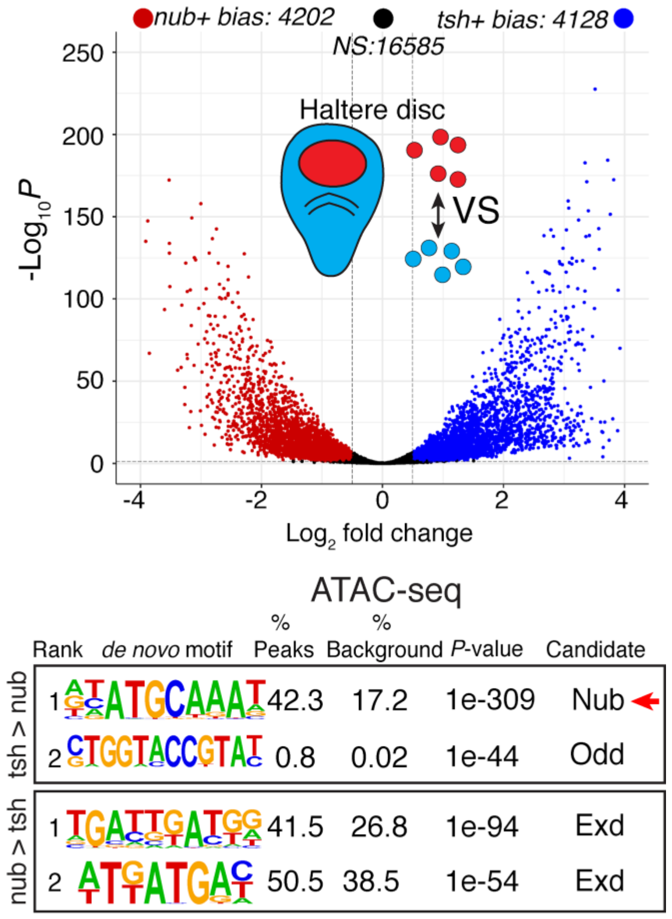


**Figure S10**

Comparison of ATAC-seq profiles from *nub*+ and *tsh*+ cell populations in the haltere imaginal disc. *de-novo* motif enrichment performed in regions with less accessibility (top) or more accessibility (bottom).

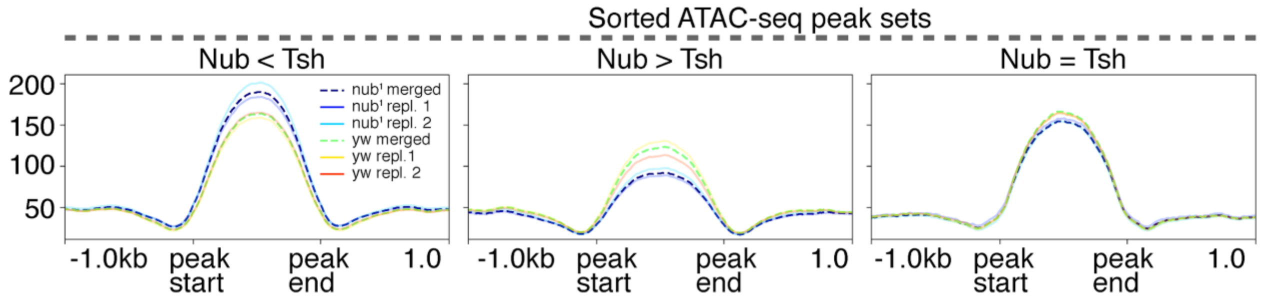


**Figure S11**

Comparison of normalized ATAC-seq read profiles from *nub¹* and wild type(*yw*) wing within genomic regions determined from sorted wild type wing discs. Dashed lines represent profiles following merging of two replicates.

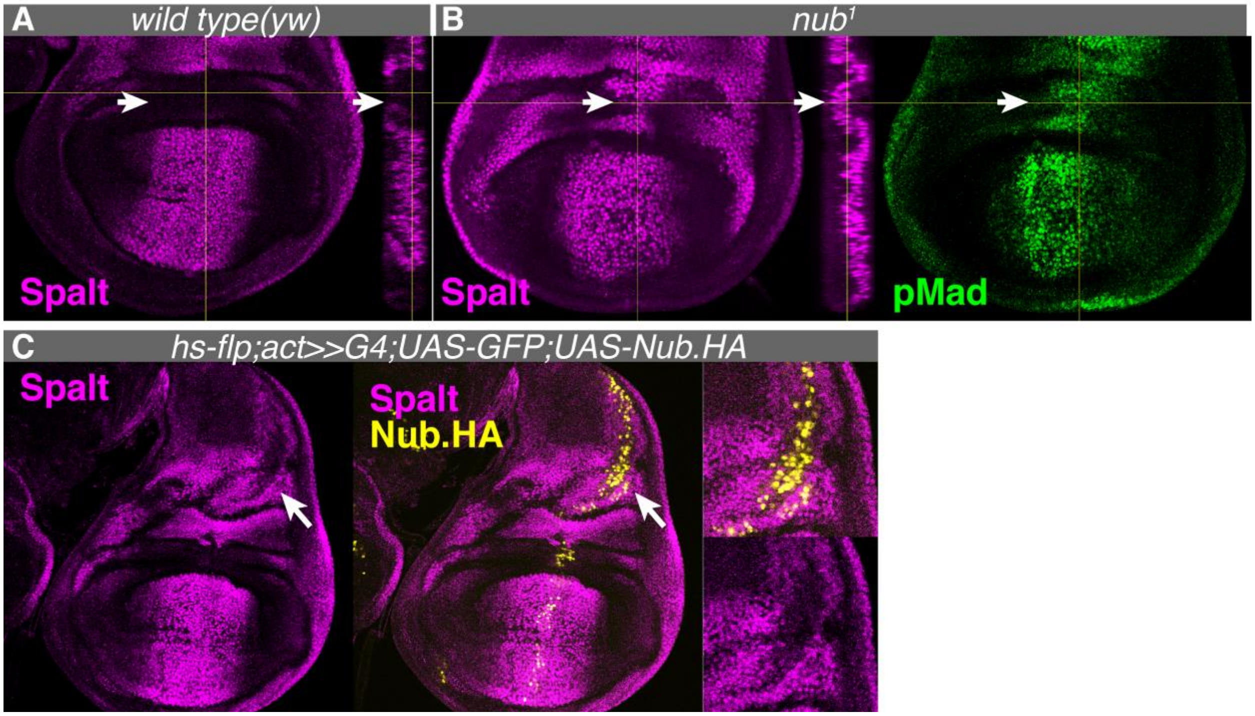


**Figure S12**

(A) Spalt expression (magenta) in wildtype wing discs. Expression is absent in the distal hinge region(arrow)

(B) Left: Spalt expression is derepressed in the distal hinge region(arrow) of *nub*$^{1}$ wing discs. Right: Dpp signalling is active in the distal hinge region where Spalt derepression is observed, indicated by phosphorylated Mad (pMad;green) antibody labeling.

(C) Ectopic expression of an HA-tagged Nub protein (yellow) in the proximal wing domain causes autonomous repression of Spalt.
