## supplemental_oligos for "A promoter-proximal silencer modifies the activity of a shared enhancer to mediate divergent expression of *nub* and *pdm2* paralogs in wing development"

| Oligos and synthesized fragments | |
|---|---|
| **Sal_tsh_1** | |
| Foward | tttttCCTGCAGGCTCTTTGCACGTATGTACATCAAATG |
| Reverse | tttttCCTAGGCATAAGAAAGCAATCCAAAAATGTCTG |
| **NPW-small deletion** | |
| gRNA#1 | TGGGCTACTGTTGGGAGACC |
| gRNA#2 | TTATTTCATAAAGCTCATAA |
| left_homology_Forward | ttttGCGGCCGCGTTTGTCTCGGGCACCTCTAC |
| left_homology_Reverse | ttttGGTACCACCCGGAGCTCTTGTCCC |
| right_homology_Forward | ttttGACGTCTGAGCTTTATGAAATAAAGACTTTATATAG |
| right_homology_Reverse | ttttACCGGTCTCGTAGGTGATATGTGTCATC |
| **NPW-large deletion** | |
| gRNA#1 | GAGGCGTTCTGAGAAGTGTT |
| gRNA#2 | TTATTTCATAAAGCTCATAA |
| left_homology_Forward | ttttGCGGCCGCCATAATAACCCTACCCGAAAGTCC |
| left_homology_Reverse | ttttGGTACCACTTCTCAGAACGCCTCAG |
| right_homology_Forward | ttttGACGTCTGAGCTTTATGAAATAAAGACTTTATATAG |
| right_homology_Reverse | ttttACCGGTCTCGTAGGTGATATGTGTCATC |
| **pdm2 promoter fragment (synthesized)** | coordinates(dm6): chr2L:12,657,539-12,657,982 |
